# NEMF-mediated CAT-tailing defines distinct branches of translocation-associated quality control

**DOI:** 10.1101/2024.08.27.610005

**Authors:** Amanda Ennis, Lihui Wang, Xiaorong Wang, Clinton Yu, Layla Saidi, Yue Xu, Sijung Yun, Lan Huang, Yihong Ye

## Abstract

Ribosome stalling during co-translational translocation at the endoplasmic reticulum (ER) causes translocon clogging and impairs ER protein biogenesis. Mammalian cells resolve translocon clogging vial a poorly characterized translocation-associated quality control (TAQC) process. Here, we combine genome-wide CRISPR screen with live cell imaging to dissect the molecular linchpin of TAQC. We show that substrates translated from mRNAs bearing a ribosome stalling poly(A) sequence are degraded by lysosomes and the proteasome, while substrates encoded by non-stop mRNAs are degraded by an unconventional ER-associated degradation (ERAD) mechanism involving ER-to-Golgi trafficking and KDEL-dependent substrate retrieval. The triaging diversity appears to result from the heterogeneity of NEMF-mediated CATylation, because a systematic characterization of representative CAT-tail mimetics establishes an AT-rich tail as a “degron” for ERAD, whereas an AG-rich tail can direct a secretory protein to the lysosome. Our study reveals an unexpected protein sorting function for CAT-tailing that safeguards ER protein biogenesis.

---

Protein homeostasis, or "proteostasis", maintenance is a critical cellular function involving various quality control mechanisms operating at the site of protein synthesis to ensure proper folding and function of nascent polypeptides. The endoplasmic reticulum (ER) is a major site of protein biogenesis, producing most secretory and membrane proteins. Accordingly, the ER is equipped with a major quality control pathway named ER-associated protein degradation (ERAD), which identifies and exports misfolded or incompletely assembled proteins from the ER to the cytoplasm for degradation by the ubiquitin-proteasome system (UPS) (Bhattacharya and Qi, 2019; Christianson and Ye, 2014; Ruggiano et al., 2014). This process involves ER chaperones for substrate recognition, ubiquitin ligase-containing membrane complexes for protein retrotranslocation, and the conserved ATPase VCP/p97, which dislocates substrates from the ER membrane to the cytosol (Vembar and Brodsky, 2008; Ye et al., 2001).

While significant progress has been made in understanding how ERAD removes misfolded proteins, less is known about the mechanism for eliminating nascent chains stalled in the Sec61 translocon due to translation arrest during co-translational protein translocation at the ER (Wang and Ye, 2020). Translation arrest and/or ribosome stalling is common during protein synthesis, triggered by defective mRNAs, mRNAs with secondary structures or difficult-to-translate sequences, such as those coding proline-rich domains of collagens (Wang et al., 2023; Yip and Shao, 2021). In the cytoplasm, a ribosome-associated quality control (RQC) pathway resolves translation stalling, starting with the ubiquitination of small ribosomal proteins (Filbeck et al., 2022; Joazeiro, 2019). Ribosome splitting factors then separate ribosomal subunits, recruiting an RQC complex consisting of the ubiquitin ligase LTN1 and NEMF (Nuclear Export Mediator Factor) (Shao et al., 2015). LTN1 ubiquitinates the nascent chain (Inada, 2020), while NEMF catalyzes CAT-tailing, adding short peptide tails to RQC substrates in an mRNA-independent manner (Howard and Frost, 2021; Kostova et al., 2017). Although the role of LTN1 in RQC is well-understood, the function of NEMF-mediated CATylation is unclear. Importantly, it is unclear whether the knowledge obtained from studying cytosolic RQC can be translated to ribosome stalling at the ER.

Ribosome stalling at the ER imposes a unique challenge to maintaining cellular proteostasis. Partially translated products are non-functional and prone to aggregation. Moreover, they can clog the Sec61 translocon since their N-termini have reached the ER lumen but their C-termini remain attached to ribosomes in the cytosol, impeding protein flow into the ER and disrupting protein biogenesis and secretion(Wang et al., 2020). Recent studies, including our own, have shown that translation stalling-induced translocon clogging activates an ER-associated ligase complex specific for the ubiquitin-like protein UFM1, resulting in the modification of the 60S ribosomal protein RPL26 with multiple UFM1 moieties (Walczak et al., 2019; Wang et al., 2020). We further showed that ribosome UFMylation activates a translocon-associated UFM1 sensor named SAYSD1, which facilitates the transfer of translocon “cloggers” to lysosomes for degradation (Wang et al., 2023; Wang and Ye, 2020). These findings define a translocation-associated quality control (TAQC) mechanism distinct from RQC. However, a contradictory study concluded that it is the proteasome that acts in conjunction with VCP/p97 to eliminate abnormal translocation products in a manner similar to cytosolic RQC(Scavone et al., 2023).

In this study, we used multiple translocon-clogging substrates, combining live cell imaging with CRISPR-mediated genetic fingerprinting, to define the fate of translocon “cloggers”. Our results reveal an unexpected role for NEMF-mediated CAT-tailing in triaging TAQC substrates, which unify previous findings into a cohesive model. We propose that TAQC is a unique protein quality control mechanism bearing features of both RQC and ERAD.

## Results

### TAQC substrates can be degraded by both lysosomes and the proteasome

To resolve the discrepancy between our studies (Wang et al., 2020; Wang et al., 2023) and that from Kopito and colleagues (Scavone et al., 2023), we used live cell fluorescence microscopy to re-analyze the fate of _SS_GFP_K20, which was shown to be a proteasome substrate (Scavone et al., 2023). _SS_GFP-K20 contains an N-terminal signal sequence (ss) for ER targeting and 20 consecutive Lys residues, encoded by a poly(A) sequence known to stall ribosome (Dimitrova et al., 2009; Juszkiewicz and Hegde, 2017). When ribosomes stall, co-translational protein translocation also pauses, generating a truncated polypeptide that clogs the translocon (Wang et al., 2020). We transfected U2OS cells with _SS_GFP_K20 and treated these cells with either a ubiquitin activating enzyme (E1) inhibitor (TAK-243), VCP/p97 inhibitor (NMS-873), lysosome inhibitor (Baf A1), proteasome inhibitor (MG132), or a combination of Baf A1 and MG132. As expected, MG132 treatment stabilized _SS_GFP_K20 in a time-dependent manner, as did NMS-873 and TAK-243 (Fig. 1A-D). Notably, MG132 and TAK-243 caused _SS_GFP_K20 to accumulate not only in the ER, but also in large nuclear puncta in ∼54% and ∼40% of the transfected cells, respectively (Fig. 1A, panels 2, 4). By contrast, NMS-873 only caused _SS_GFP_K20 to accumulate in the ER (panel 5). Co-staining with a nucleolin antibody confirmed the nuclear puncta as nucleoli (Fig. 1E), suggesting that _SS_GFP_K20 is a retrotranslocation substrate. When its degradation in the cytosol is inhibited, _SS_GFP_K20 was translocated into nucleoli, likely because it contains a Lys-rich nucleolar localization sequence (Martin et al., 2015). Surprisingly, Baf A1 treatment also stabilized _SS_GFP_K20, as ∼28% of the transfected cells showed vesicle-like puncta, ∼22% cells contained nucleolus-like puncta, and ∼5% cells had both (Fig. 1A, panel 3). Co-immunostaining and live cell confocal imaging confirmed extensive co-localization of _SS_GFP_K20 with either endogenous lysosomal protein LAMP1 or transfected LAMP1-mCherry (Fig. 1F, Fig. S1A). Although it is unclear why Baf A1 caused _SS_GFP_K20 to accumulate in nucleoli in some cells, the observed lysosomal accumulation suggests that _SS_GFP_K20 can also be degraded by lysosomes . Interestingly, in cells co-treated with MG132 and Baf A1, we observed nucleolar and lysosomal accumulation of _SS_GFP_K20 in distinct populations of the cells (∼34% each); only a small number of the cells (∼6%) had both patterns (Fig. 1A, panel 6). These results suggest that most cells use either the proteasome or lysosome to degrade _SS_GFP_K20 (see discussions).

**Figure 1.**
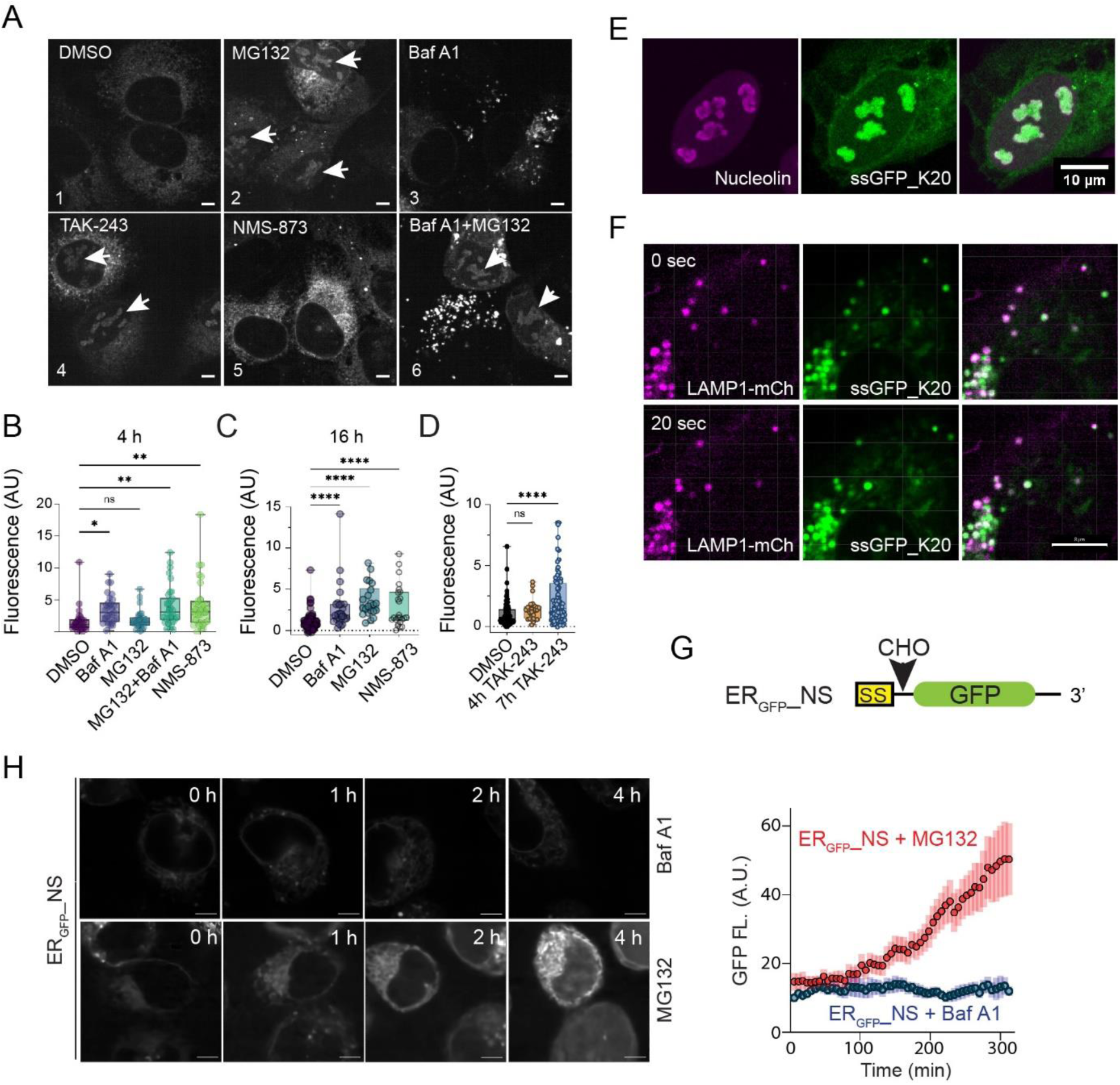
TAQC substrates can be degraded by both the proteasome and lysosomes. (A) U2OS cells transfected with ssGFP_K20 were treated with the indicated inhibitors for 4 h and imaged. Arrowheads indicate nucleolar localization of _SS_GFP_K20. Scale bars, 5 μm. (B-D) Quantification of GFP fluorescence in _SS_GFP_K20-transfected cells treated with the indicated inhibitors. Shown are box and whiskers plot with all points. The boxes show the Upper Quartile, the Median and the Lower Quartile. ns, not significant; *, p<0.05; **, p<0.01; ****, p<0.001 by one-way ANOVA’s Dunnett’s multiple comparison test. n > 20 cells in randomly selected field from two independent experiments. (E) _SS_GFP_K20 accumulates in nucleoli in MG132-treated cells. U2OS cells expressing _SS_GFP_K20 were treated with MG132 and stained with anti-nucleolin antibodies (magenta). Scale bar, 10 μm. (F) _SS_GFP_K20 accumulates in LAMP1-mCherry (mCh) positive lysosomes in Baf A1-treated cells. Shown are two time points from a live cell imaging experiment. Scale bar, 8 μm. (G) A schematic diagram of the ER_GFP__NS reporter. (H) Left panels, ER_GFP__NS-transfected HEK293T cells were treated with Baf A1 (200 nM) or MG132 (20 µM) and then imaged continuously for 6 h. The graph shows the quantification of the experiment. Error bars, s.e.m. n=6 cells for each condition.

To further dissect the degradation mechanism of TAQC, we used an additional ss-containing ribosome stalling reporter, ER_GFP__NS. The mRNA encoding ER_GFP__NS lacks any stop codons for translation termination, causing ribosome stalling (Fig. 1G). When we treated HEK293T cells stably expressing ER_GFP__NS with Baf A1 or MG132, live cell confocal microscopy showed that MG132, but not Baf A1, increased GFP fluorescence over time (Fig. 1H). Pulse chase experiments confirmed that ER_GFP__NS was short-lived, and its turnover was inhibited by MG132, but not by lysosomal inhibitors chloroquine or Baf A1 (Fig. S1B, C). Collectively, these results strongly indicate that ER_GFP__NS degradation is entirely mediated by the proteasome.

### ER_GFP__NS is degraded by an ERAD-like mechanism involving SAYSD1 and the TRAPPC complex

The finding that ER_GFP__NS’s degradation is entirely mediated by the proteasome is surprising because like ER_GFP__K20 and _SS_GFP_K20, the degradation of ER_GFP__NS was known to involve UFM1 (Wang et al., 2023). We therefore combined cell fractionation with inhibitor treatment and *UFM1* knockdown to examine how UFM1 acts with the proteasome in ER_GFP__NS degradation. Consistent with UFM1 being a positive TAQC facilitator, immunoblotting analysis of whole cell extracts detected ER_GFP__NS accumulation in UFM1-depleted cells, but it appeared in a full-length (∼45kDa) and an SDS-resistant high molecular weight (HMW) form (Fig. 2A, lane 2). The HMW ER_GFP__NS “smear” was not obvious in proteasome inhibitor-treated cells, excluding ubiquitination as the main cause of this molecular weight shift. By contrast, MG132 treatment caused accumulation of mostly full-length and a cleaved ER_GFP__NS species, each containing a fast- and a slow-migrating band on the blot (lane 3 vs. 1). Biochemical fractionation showed that ER-enriched microsomes from MG132-treated cells had all four ER_GFP__NS bands, but the corresponding cytosolic faction only contained the fast-migrating band of each species (Fig. 2B, lane 4 vs. 3). When the membrane fractions from MG132-exposed cells were treated with Endo H, the slow-migrating ER_GFP__NS forms were converted to the corresponding fast-migrating species (Lane 6 vs 5), suggesting that the slow-migrating ER_GFP__NS band was caused by N-glycosylation. Collectively, these results suggest that ER_GFP__NS is retrotranslocated from the ER and deglycosylated, similarly to previously reported glycol-ERAD substrates (Shamu et al., 1999). Consistent with this notion, the VCP/p97 inhibitor NMS-873 only caused ER_GFP__NS to accumulate in the glycosylated forms (Fig. 2A, lane 5). Knockdown of UFM1 in MG132-or NMS0873-treated cells reduced both deglycosylated and cleaved ER_GFP__NS (Fig. 2A, lane 4 vs. 3), suggesting that UFMylation acts upstream of retrotranslocation.

**Figure 2.**
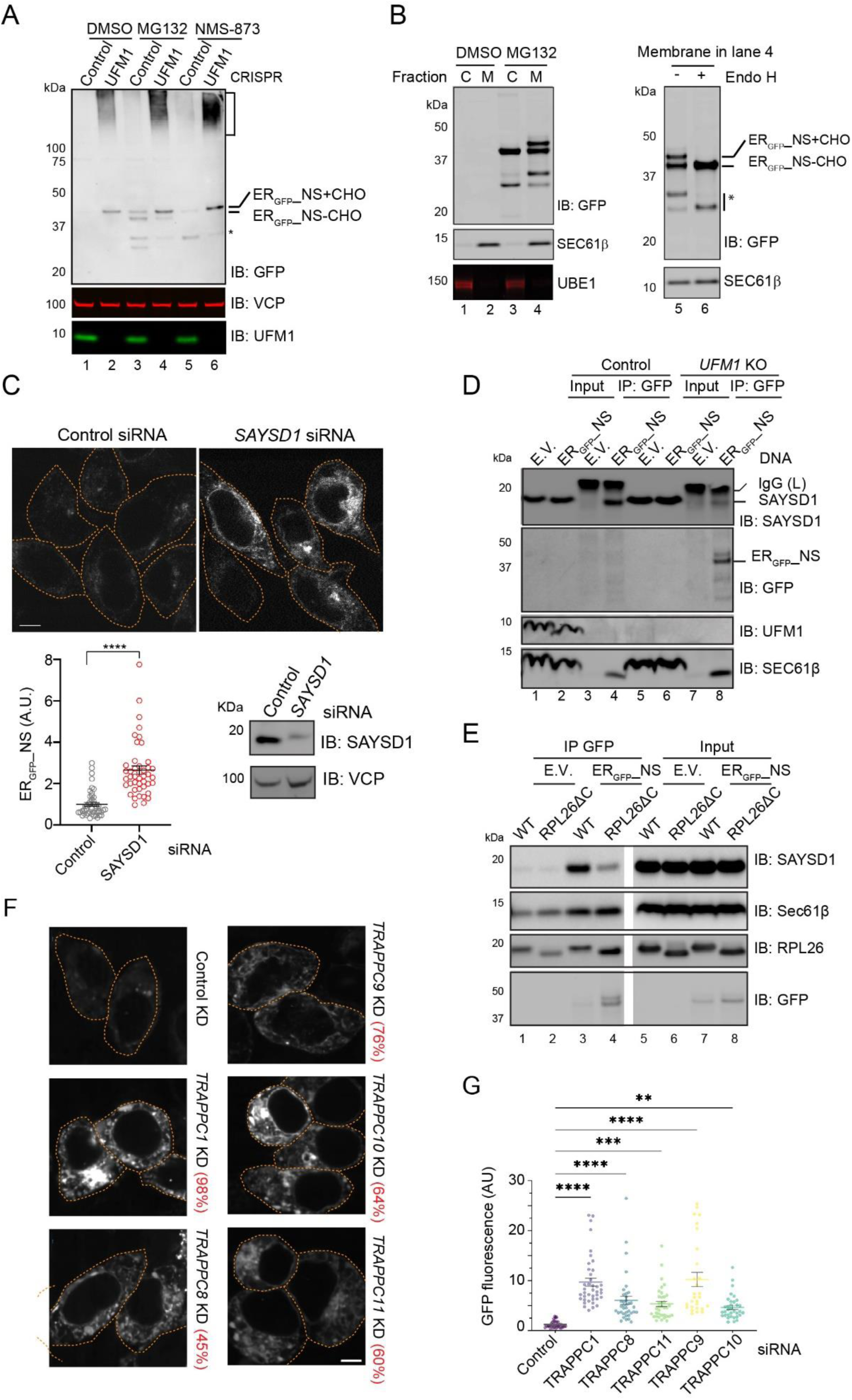
Ribosome UFMylation, SAYSD1 and TRAPPC complex facilitates ER_GFP__NS degradation via an ERAD-like mechanism. (A) Control or UFM1 knockout HEK293T cells were transfected with ER_GFP__NS and treated with the indicated inhibitors for 6 h. Cell extracts were analyzed by immunoblotting (IB). +CHO, glycosylated; - CHO, deglycosylated. (B) ER_GFP__NS-expressing cells treated as indicated were fractionated into a membrane (M) and cytosol (C) fraction before immunoblotting (left panel). Right panel, the membrane fraction used in lane 4 was treated with Endo H and then analyzed by immunoblotting. *, a cleaved ER_GFP__NS fragment. (C) Top panels, HEK293T cells were transfected with the indicated siRNA together with ER_GFP__NS. Cells were imaged 48 h post transfection. A fraction of the cells was analyzed by immunoblotting to confirm the knockdown efficiency (bottom right). The graph shows the quantification results from three independent experiment. A.U., arbitrary unit. N>40 cells. ****, p<0.0001 by unpaired Student’s t-test. (D) Control or UFM1 KO cells were transfected with either empty vector (EV) or ER_GFP__NS. Cells were lysed in CHAPS buffer. The lysates were subject to immunoprecipitation with GFP antibodies before immunoblotting. (E) As in D, except that RPL26dC and an isogenic parent wildtype (WT) line were used. (F) HEK293T cells transfected with the indicated siRNA and ER_GFP__NS were analyzed by confocal microscopy. The numbers indicate average knockdown efficiency determined by qRT-PCR from three independent experiments. (G) Quantification of experiments represented in (F). At least 30 randomly selected cells from two independent experiments were analyzed. Error bars, s.e.m. **, p<0.01, ***, p<0.001, ****, p<0.0001 by one-way ANOVA’s Dunnett’s multiple comparison test.

Since our previous study demonstrated the involvement of a translocon-associated, ribosome- and UFM1-binding protein SAYSD1, and the TRAPPC complex in the ER-to-Golgi trafficking and lysosomal degradation of ER_GFP__K20 (Wang et al., 2023), we tested whether these factors also facilitate ER_GFP__NS turnover.

To test whether SAYSD1 is involved in ER_GFP__NS turnover, we used *SAYSD1* specific siRNA to knock down its expression in ER_GFP__NS-expressing HEK293T cells. Confocal microscopy detected a ∼3-fold increase in ER_GFP__NS fluorescence after *SAYSD1* depletion (Fig. 2C). Co-immunostaining showed that ER_GFP__NS accumulated in *SAYSD1* knockdown cells was largely co-localized with the ER marker calreticulin (Fig. S2A). We next tested whether ER_GFP__NS interacts with SAYSD1 by co-immunoprecipitation. To this end, we transfected wildtype (WT) and *UFM1* knockout cells with either an empty vector or an ER_GFP__NS-expressing construct. Immunoprecipitation with GFP antibodies pulled down ER_GFP__NS together with endogenous SAYSD1 and Sec61β from WT cells (Fig. 2D, lane 4). By contrast, in *UFM1* knockout cells, although more ER_GFP__NS and Sec61 β were in precipitated samples, less SAYSD1 was co-precipitated (lane 8 vs. 4). This result demonstrates an UFMylation dependent interaction between ER_GFP__NS and SAYSD1, similarly to what was reported for ER_GFP__K20 (Wang et al., 2023).

To determine whether the interaction with SAYSD1 depends on UFMylation of RPL26, we repeated the co-immunoprecipitation experiment using a CRISPR-engineered knock-in cell line that has the RPL26 C-terminal UFMylation sites removed (RPL25ΔC) (Wang et al., 2023). Like in *UFM1* knockout cells, GFP antibodies precipitated more ER_GFP__NS from RPL26ΔC cells, but less SAYSD1 was co-precipitated (Fig. 2E, lane 4 vs. 3), suggesting that RPL26 UFMylation enhances the interaction of ER_GFP_-NS with SAYSD1. By contract, the interactions of ER_GFP__NS with the Sec61 translocon and the ribosome were not affected in RPL26ΔC cells. Together, these findings implicate SAYSD1 as a positive regulator, engaging ER_GFP__NS in a RPL26 UFMylation-dependent manner to facilitate its degradation.

To test whether ER_GFP__NS’s degradation also requires the TRAPPC complex, we used siRNA to knock down several components of the TRAPPC complex in ER_GFP__NS cells. Quantitative RT-PCR confirmed the knockdown efficiency, ranging from 45% to 98% (Fig. 2F). Fluorescence confocal microscopy showed significant upregulation of ER_GFP__NS in cells depleted of TRAPPC1, TRAPPC8, TRAPPC9, TRAPPC10, or TRAPPC11 (Fig. 2F, G). Co-immunostaining showed that in cells depleted of TRAPPC8, ER_GFP__NS mainly accumulated in a peri-nuclear region overlapping with the ER marker calreticulin (Fig. S2B). Thus, TRAPPC-mediated ER-to-Golgi trafficking also facilitates ER_GFP__NS degradation.

### ER_GFP__NS degradation requires the ER-retrieval pathway and ERAD

To obtain the genetic fingerprint of ER_GFP__NS degradation, we conducted an unbiased genome-wide CRISPR screen using HEK293T cells stably expressing ER_GFP__NS. To this end, we targeted the ∼20,000 human genes each with six single guide RNAs (sgRNAs), expressed from a lentivirus-based CRISPR-Cas9 library (Sanjana et al., 2014). The transduced cells were separated into a GFP-high (∼10%) and the remaining (∼90%) population by flow cytometry. We then used PCR amplification and high-throughput sequencing to identify sgDNAs enriched in the GFP high population. Statistical analysis revealed 824 genes whose inactivation elevated ER_GFP__NS (Fig. 3A). We compared this list to the genes identified previously in a similar screen using ER_GFP__K20 as the reporter (Wang et al., 2023). Since these two substrates are degraded by different mechanisms, most of the identified genes did not overlap. Nevertheless, 88 genes (∼5%), when inactivated, increased GFP fluorescence of both ER_GFP__K20 and ER_GFP__NS. STRING-based protein network analyses showed that most common “hits” were associated with RNA catabolism, which likely affected the reporters via mRNA abundance. Notably, the overlapping gene list did include several UFMylation components and a few ER-to-Golgi trafficking regulators, such as the TRAPPC complex components (Fig. 3B, Fig. S3), further validating the involvement of UFMylation and ER-to-Golgi trafficking in ER_GFP__NS degradation. Our screen failed to uncover SAYSD1 and some TRAPPC complex components, suggesting that the screen is not saturated.

**Figure 3.**
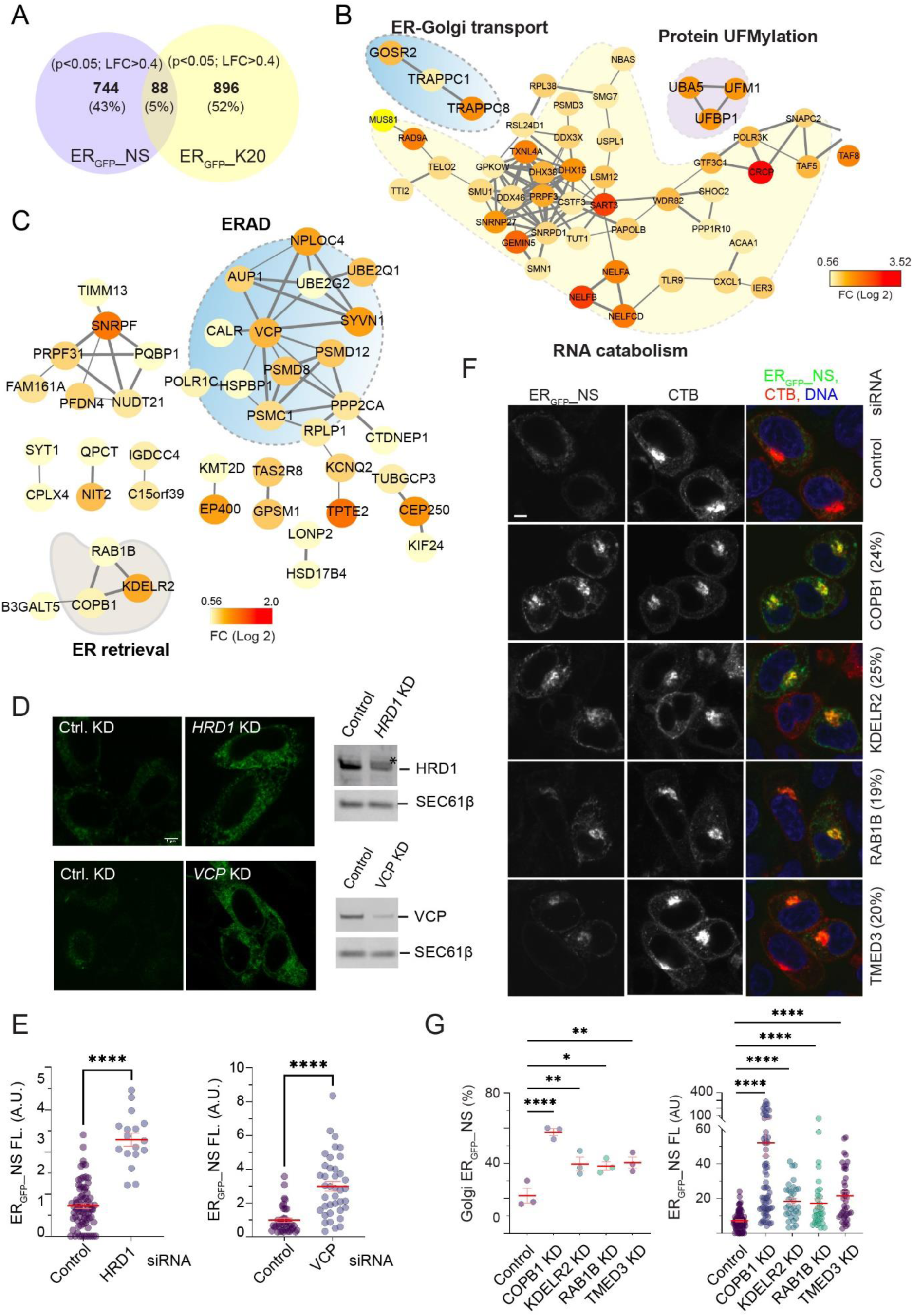
A CRISPR screen reveals the genetic fingerprint of ER_GFP__NS degradation. (A) A Venn diagram shows the number of genes, when inactivated by CRISPR, increased ER_GFP__NS and ER_GFP__K20, respectively. LFC, log fold change. (B) A STRING network analysis reveals gene networks that affect the steady state levels for both ER_GFP__NS and ER_GFP__K20. (C) STRING networks affecting only the level of ER_GFP__NS. Node color in (B, C) indicates fold enrichment in GFP high populations. (D) HEK293T cells were transfected with control, Hrd1, or VCP/p97 siRNA together with ER_GFP__NS. Cells were imaged by fluorescence confocal microscopy to measure ER_GFP__NS levels (left) or by immunoblotting (right) to confirm gene knockdown. Scale bar, 5 μm. (E) Quantification of two independent experiments as shown in (D). Error bars, s.e.m., ****, p<0.0001 by unpaired Student’s t-test. (F) HEK293T cells were transfected with the indicated siRNA followed by ER_GFP__NS. Cells were stained by Alexa^594^-labeled cholera toxin subunit B to identify the Golgi. (G) Quantification of percentage transfected cells showing a Golgi localization of ER_GFP__NS (left) or the total fluorescence intensity of ER_GFP__NS (right) from three independent experiments. Error bars, s.e.m., *, p<0.05; **, p<0.01, ****, p<0.0001 by one-way ANOVA’s Dunnett’s multiple comparison test.

Next, we used STRING-based analysis to identify functional networks associated only with ER_GFP__NS but not ER_GFP__K20 degradation. This approach identified one major network ‒ ERAD and several smaller ones (Fig. 3C). While the role of these small networks in ER_GFP__NS regulation awaits further characterization, the identification of ERAD as a major regulator of ER_GFP__NS turnover, combined with the observed sensitivity of this substrate to proteasome and VCP inhibitors, suggested ERAD as a critical mediator of ER_GFP__NS degradation. Indeed, siRNA-mediated knockdown of Hrd1/SYVN1 or VCP/p97, two major ERAD genes identified by our CRISPR screen, significantly increased ER_GFP__NS abundance in the cell (Fig. 3D, E).

At first glance, the degradation of ER_GFP__NS by ERAD appears inconsistent with the involvement of UFMylation, SAYSD1, and TRAPPC-dependent ER-to-Golgi trafficking in this pathway. However, further examination of the gene networks specific for ER_GFP_-NS identified several components of the KDEL-dependent ER-retrieval pathway, including KDELR2, COPB1, and RAB1B (Fig. 3C), suggesting that ER_GFP__NS might be exported to the Golgi and then retrieved back to the ER before degradation by ERAD. Consistent with this view, when we used siRNAs to knock down the identified ER retrieval genes in ER_GFP__NS cells, we found that knockdown of KDELR2, COPB1, RAB1B, and TMED3 not only increased ER_GFP__NS fluorescence, but also caused its accumulation in a peri-nuclear region, co-localizing with the Golgi marker, cholera toxin B (CTB) (Fig. 3F, G). Collectively, these results suggest that following translation stalling and ribosome UFMylation, ER_GFP__NS uses a TRAPPC-dependent mechanism to reach the Golgi and is then retrieved back to the ER before undergoing retrotranslocation and ERAD (see discussion).

### LTN1 and NEMF regulate ER_GFP__NS degradation

Previous studies showed that the degradation of ER_GFP__K20 and _SS_GFP_K20 involves LTN1 and NEMF (Scavone et al., 2023; Wang et al., 2023). These proteins are known to function as an RQC complex: LTN1 catalyzes ubiquitination while NEMF mediates CATylation of translation-stalled proteins (Inada, 2020). To explore whether these functions are conserved in TAQC, we transfected U2OS cells with ER_GFP__NS and siRNAs targeting *LTN1* or *NEMF*. Confocal fluorescent microscopy showed that knockdown of NEMF or LTN1 caused a ∼5-fold upregulation of ER_GFP__NS in the cell (Fig. 4A-C). Immunoblotting further confirmed that knockdown of *LTN1* or *NEMF* increased the steady state level of ER_GFP__NS, similarly to knockdown of UFM1; this phenotype was observed regardless of whether knockdown cells were treated with a low concentration of anisomycin (ANS), which induces ribosome stalling and RPL26 UFMylation (Fig. 4D, E). Noticeably, *NEMF* or *LTN1* knockdown did not cause ER_GFP__NS to accumulate in the SDS-resistant HMW “smear” as seen in *UFM1* knockdown cells (Fig. 4D, lanes 3, 4 vs. lanes 5-8). However, when NEMF and UFM1 were knocked down simultaneously, no HMW “smear” was observed (Fig. 4D, lanes 9, 10 vs. 3, 4; Fig. 4F). Given the well-established function of NEMF in the CATylation of translation-stalled polypeptides, the HMW ER_GFP__NS species likely results from NEMF-mediated CATylation.

**Figure 4.**
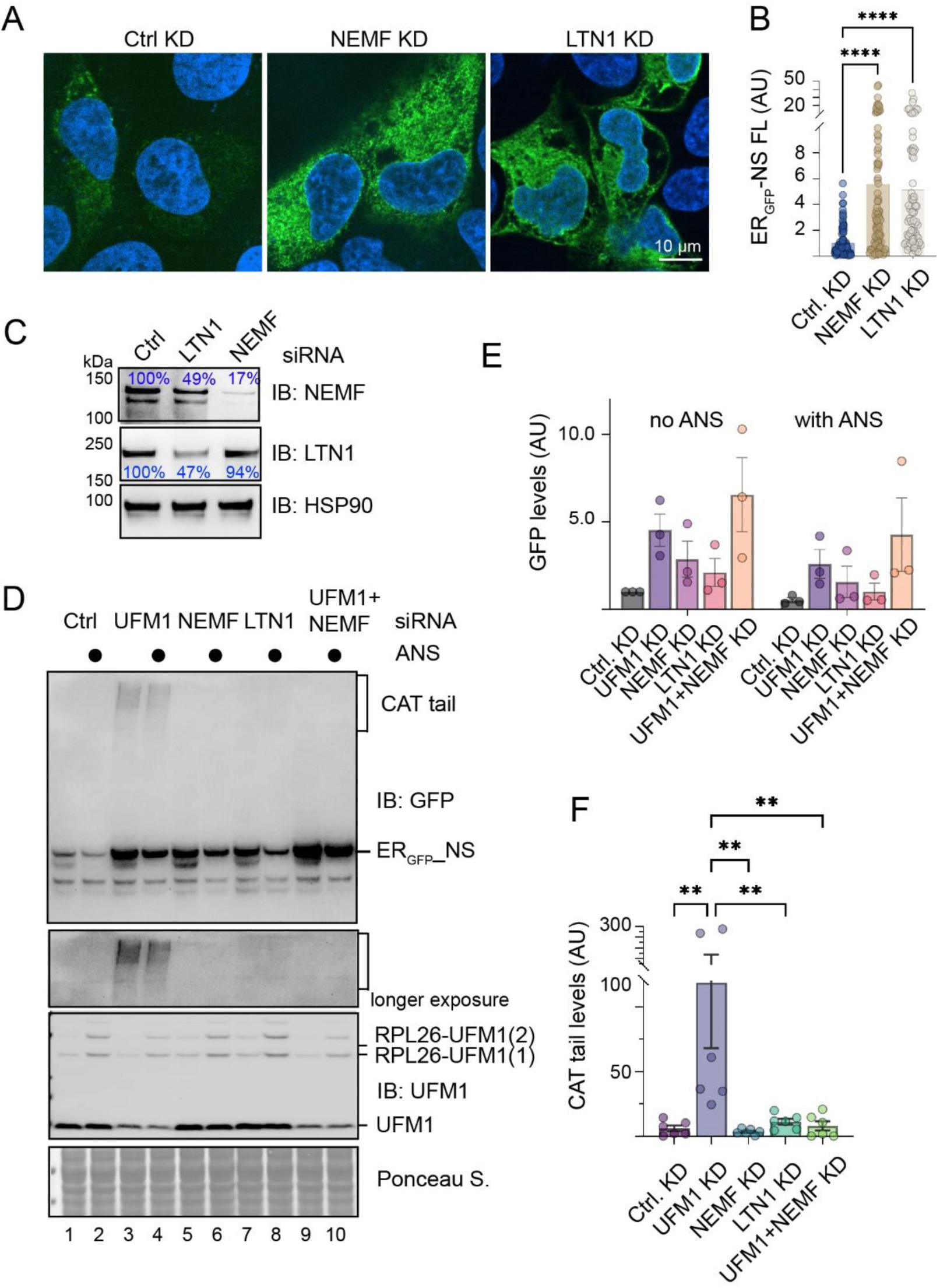
LTN1 and NEMF1 facilitates ER_GFP__NS degradation. (A) U2OS cells transfected with the indicated siRNA and ER_GFP__NS were stained with Hoechst and imaged. (B) Quantification of 2 independent experiments as shown in A. Error bars, s.e.m., ****, p0.0001 by one-way ANOVA’s Dunnett’s multiple comparison test. (C) A fraction of transfected cells in A were analyzed by immunoblotting to confirm knockdown efficiency. (D) HEK293E cells transfected with the indicated siRNA and ER_GFP__NS were treated with anisomycin (ANS, 200nM) for 1 h. Cell lysates were analyzed by immunoblotting. (E) Quantification of ER_GFP__NS levels in three independent experiments represented by D. Error bars, s.e.m. (F) Quantification of HMW “smear” in the indicated knockdown cells. Error bars, s.e.m., **, p<0.01 by one-way ANOVA’s Dunnett’s multiple comparison test.

### CAT-tailing alters the trafficking path of a secretory protein

Since CATylation usually appends a small tag of less than 10 kDa to stalled nascent chains as reported for _SS_GFP_K20 and RQC substrates (Mizuno et al., 2021; Scavone et al., 2023), the large molecular weight shift seen with ER_GFP__NS CATylation suggested that the tag appended to ER_GFP__NS might be more prone to aggregation than that on _SS_GFP_K20. We therefore postulated that the variability in tag sequence might account for the different degradation mechanisms. To test this hypothesis, we appended a set of artificial CAT-tails to the C-terminus of a signal sequence-bearing model secretory protein (GFP_ER_) to see whether these tags could affect GFP_ER_ trafficking. Since CAT-tails formed in mammalian cells are enriched with amino acids Ala, Thr, and Gly, with Ala being the predominant one (Udagawa et al., 2021), we added either 10 or 20 AT repeats or 10 AG repeats to the C-terminus of GFP_ER_. We also added 5 Lys residues before these repeats because this is the average number of Lys residues translated by ribosomes after they encounter a poly(A) sequence (Udagawa et al., 2021). Additionally, we included constructs containing either 15 or 30 Ala residues (Fig. 5A). This strategy bypassed translation stalling and NEMF-mediated CATylation to create a collection of “CAT-tail mimetics.” As controls, we used either GFP_ER,_ which lacks any CAT-tails, or GFP_ER-ext_, which contains an irrelevant C-terminal extension of similar size as those on “CAT tail mimetics.”

**Figure 5.**
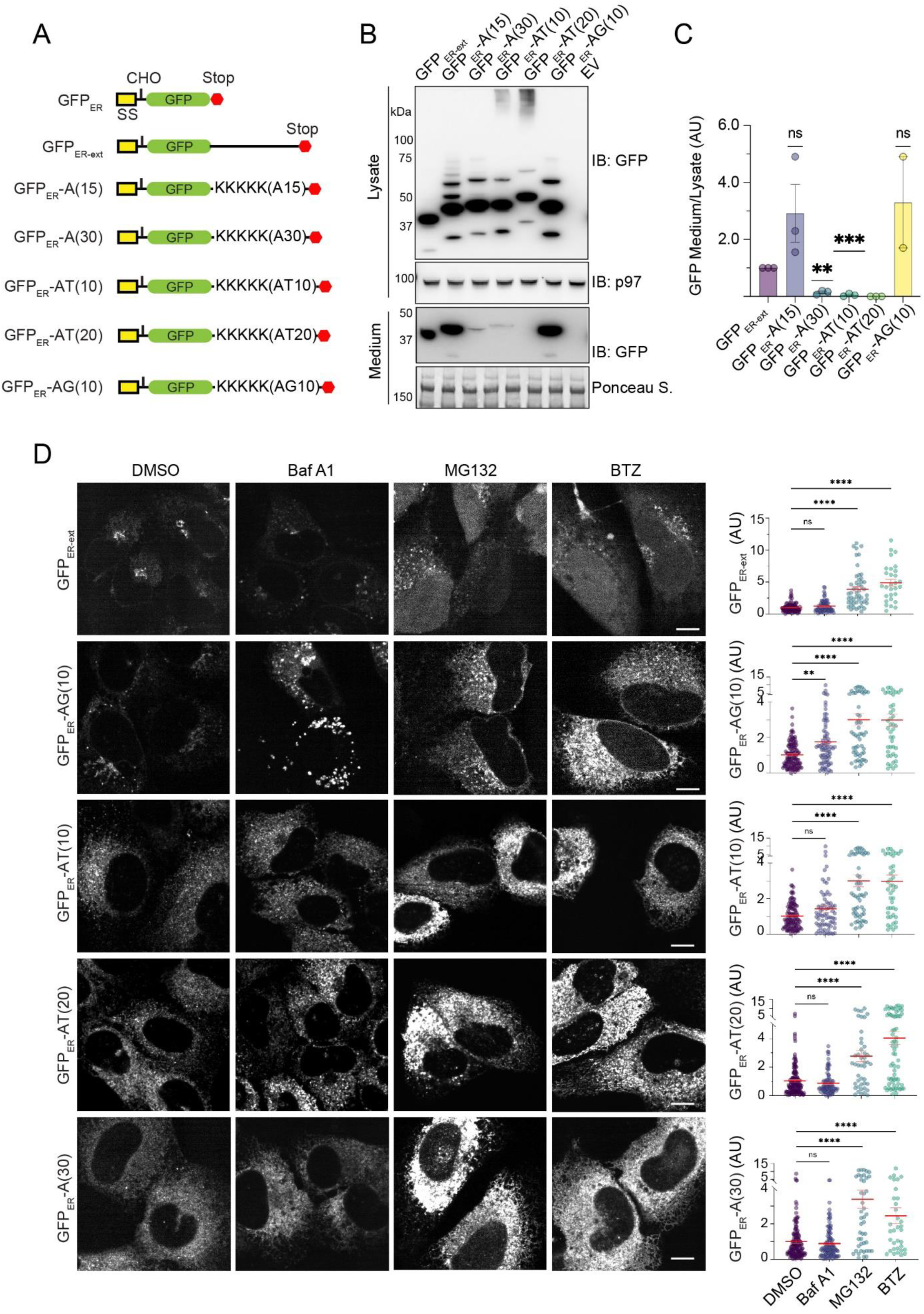
Distinct triaging options for CAT tail mimetics. (A) A schematic illustration of CAT tail mimetics and controls. (B) Cell lysate and conditioned medium from HEK293T cells transfected with the indicated constructs were analyzed by immunoblotting. EV, empty vector. (C) Quantification of the experiments as shown in B. Error bars, s.e.m., n=3. ns, not significant. **, p<0.01, ***, p<0.001 by unpaired Student’s t-test. (D) U2OS cells transfected with the CAT tail mimetics and controls were treated with the indicated inhibitors for 6 h and imaged. BTZ, Bortezomib. The graphs show the quantification of GFP fluorescence from two independent experiments. Error bars, s.e.m., *, p<0.05; **, p<0.01, ****, p<0.0001 by one-way ANOVA’s Dunnett’s multiple comparison test.

We first measured the secretion efficiency of these “CAT-tail mimetics.” We transfected these constructs into HEK293T cells and used immunoblotting to measure the secreted GFP_ER_ variants in conditioned medium, which were normalized to the corresponding protein in the cell lysate. As expected, both GFP_ER_ and GFP_ER-ext_ were readily secreted (Fig. 5B, C, Fig. S4A), and treating cells with Brefeldin A, a drug known to disrupt the Golgi structure, abolished GFP_ER_ secretion whereas MG132 had no effect (Fig. S4A). The secretion of GFP_ER_-A(15) and GFP_ER_-AG(10) was comparable to that of GFP_ER-ext_ (Fig. 5B, C). By contrast, the secretion of GFP_ER_-A(30), GFP_ER_-AT(10), and GFP_ER_-AT(20) was significantly abrogated compared to GFP_ER_ or GFP_ER-ext_ (Fig. 5B, C, Fig. S4A), suggesting that A(30), AT(10), and AT(20) tags effectively retain GFP_ER_ in the cell. Interestingly, immunoblotting also detected a fraction of GFP_ER_-AT(10) and GFP_ER_-AT(20) in HMW “smear” reminiscent of ER_GFP__NS in *UFM1* knockdown cells (Fig. 5B). Thus, the CAT-tail formed on ER_GFP__NS appears similar to AT(10) or AT(20) in their aggregation propensity.

We next used confocal fluorescent microscopy to test whether these “CAT-tail mimetics” were degraded in cells. We transfected these “CAT-tail mimetics” or GFP_ER-ext_ as a control into U2OS cells and treated the cells with either Baf A1 or two different proteasome inhibitors: MG132 and Bortezomib (BTZ). Consistent with the observed secretion of GFP_ER-ext_ and GFP_ER_-AG(10), these proteins were less abundant in cells than GFP_ER_-AT(10), GFP_ER_-AT(20), and GFP_ER_-A(30). After inhibitor treatment, the fluorescence intensity of GFP_ER-ext_ was modestly increased by either MG132 or Bortezomib, but not by Baf A1 (Fig. 5D). Since GFP_ER-ext_ accumulation could be detected in the nucleus, it appears that a fraction of GFP_ER-ext_ is retained in the ER and degraded by ERAD, probably due to a folding defect caused by either protein overexpression or the appended tag. Compared to GFP_ER-ext_, GFP_ER_-AT(10), GFP_ER_-AT(20), and GFP_ER_-A(30) were accumulated to much higher levels in MG132- and Bortezomib-treated cells. Since Baf A1 treatment did not affect their levels (Fig. 5D), we concluded that the AT(10), AT(20), and A(30) tails effectively target GFP_ER_ for degradation by the proteasome. By contrast, the GFP_ER_-AG(10) level was increased in cells treated with either Baf A1, MG132, or Bortezomib (Fig. 4D), suggesting that GFP_ER_-AG(10) can be degraded by either the lysosome or proteasome.

### CAT-tails can serve as a triaging signal

To understand how CAT-tailing differentially impacts TAQC, we focused our analysis on AT(10) and AG(10) because these two tails caused drastically different phenotypes on GFP_ER_ trafficking despite minimum sequence divergence: AT(10) inhibited GFP_ER_ secretion, targeted it for proteasomal degradation, and caused some GFP_ER_ to aggregate, while AG(10) had minimal effect on GFP_ER_ secretion, but somehow targeted a fraction of GFP_ER_ for lysosomal degradation.

We first used the RUSH assay to better define the trafficking path of GFP_ER_-AG(10) (Boncompain et al., 2012). To this end, we added a streptavidin-binding peptide (SBP) to GFP_ER_-AG(10) to trap a fraction of GFP_ER_-AG(10) at the ER by an ER-localized Streptavidin “hook”, Ii-Str (Fig. 6A) (Boncompain et al., 2012). After addition of biotin to release GFP_ER_-SBP-AG(10) from the “hook”, live cell imaging detected the movement of GFP_ER_-SBP-AG(10) to a peri-nuclear region and the formation of some GFP_ER_-SBP-AG(10)-containing vesicles (Fig. 6B, top panels, Video 1). As expect, this phenotype was not seen without biotin addition or in cells lacking the Ii-Str “hook” (Fig. S4B). When cells were pre-treated with Baf A1, we detected more GFP_ER_-SBP-AG(10) in the peri-nuclear region at time zero, forming a typical horseshoe-shaped Golgi pattern. After biotin treatment, more GFP_ER_-SBP-AG(10) was detected in vesicles (bottom panels). Many GFP_ER_-AG(10)-containing vesicles were LAMP1 positive, confirming their identity as late endosomes or lysosomes (Fig. 6C). These results suggest that GFP_ER_-AG(10) is transported from the ER to Golgi, but after arrival at the Golgi, some GFP_ER_-AG(10) was diverted to lysosomes. Thus, AG(10) appears to have a lysosome targeting function.

**Figure 6.**
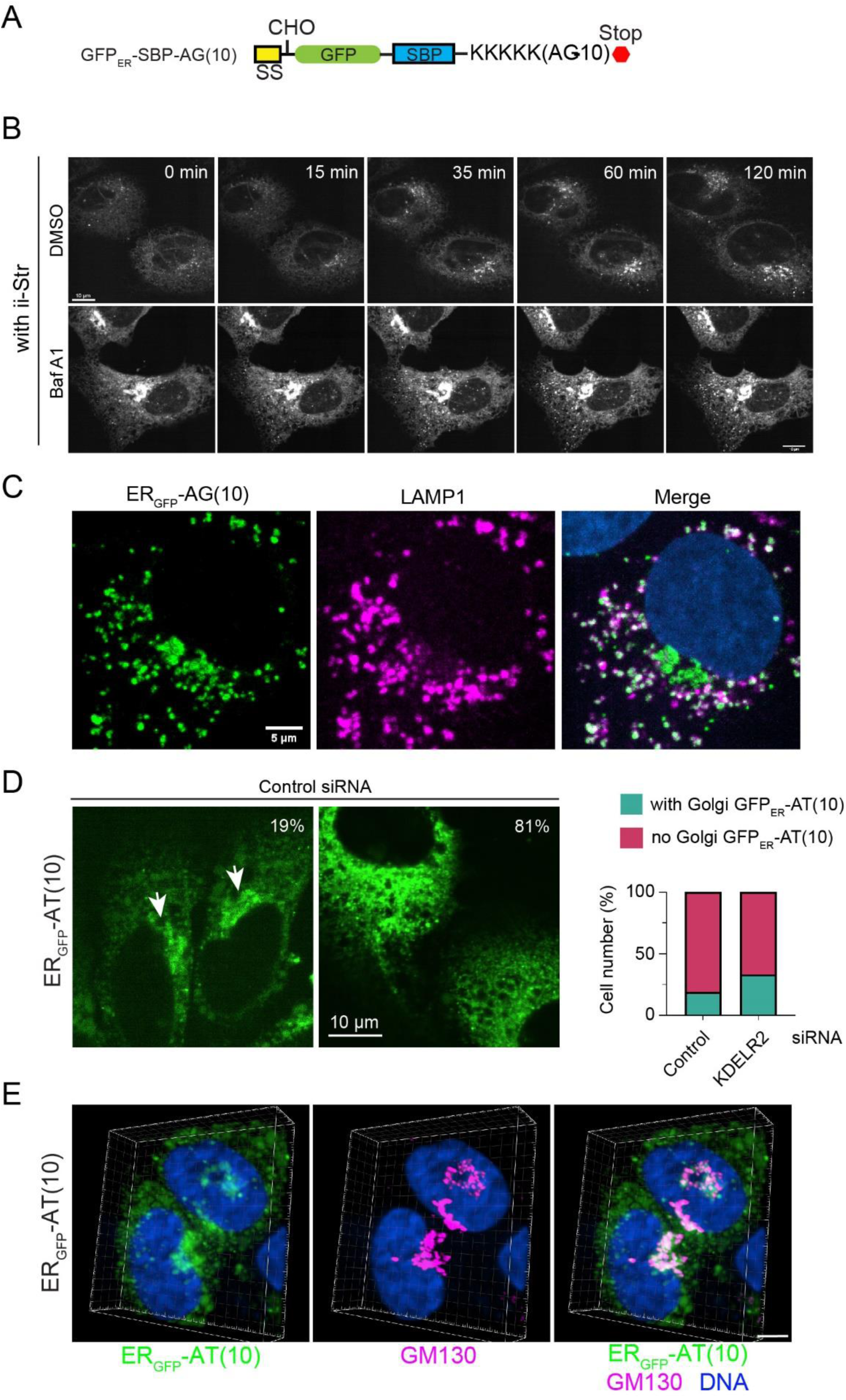
CAT tail mimetics are sorted at the Golgi. (A) A schematic diagram of the construct used in the RUSH assay. (B) Selected frames from a representative RUSH assay using U2OS cells transfected with ER_GFP_-SBP-AG(10) and Ii-Str. Where indicated, cells were pre-treated with Baf A1 (200 nM) for 2 h before imaging. Scale bars, 10 μm. (C) U2OS cells transfected with ER_GFP_-AG(10) were stained with LAMP1 antibodies (magenta) and Hoechst (blue). Scale bar, 5 μm. (D) U2OS cells transfected with ER_GFP_-AT(10) and a control siRNA were imaged and scored. Arrows indicated Golgi localization of ER_GFP_-AT(10) in a fraction of cells. The graph shows the quantification of two independent experiments. Scale bar, 10 μm. (E) U2OS cells transfected with ER_GFP_-AT(10) were stained with a GM130 antibody (magenta) and Hoechst (blue). 3D confocal imaging was used to reconstruct the localization of ER_GFP_-AT(10) in the peri-nuclear region. Scale bar, 5 μm.

For AT(10) tail, several lines of evidence suggest that this tag does not serve merely as a degron to degrade GFP_ER_ at the ER. Firstly, when AT(10) was attached to cytosolic GFP, it did not effectively target GFPcyto for proteasomal degradation (Fig. S4C). Importantly, although GFP_ER_-AT(10) was mainly localized in the ER under both basal and MG132-treated conditions, we detected GFP_ER_-AT(10) in a peri-nuclear region in ∼19% untreated cells (Fig. 6D). The morphology of the peri-nuclear GFP_ER_-AT(10)-containing structure suggested it as the Golgi. Indeed, two-color fluorescence 3D imaging showed extensive colocalization of peri-nuclear GFP_ER_-AT(10) with the Golgi marker GM130 (Fig. 6E, Video 2). Interestingly, when GFP_ER_-AT(10)-expressing cells were transfected with KDELR2 siRNA to inhibit the ER-retrieval pathway, Golgi-localized GFP_ER_-AT(10) was seen in ∼33% of the cells (Fig. 6D). Despite the small increase in the percentage of cells bearing Golgi-localized GFP_ER_-AT(10), KDELR2 knockdown did not significantly increase the GFP_ER_-AT(10) level (Fig. S5), probably because only a fraction of GFP_ER_-AT(10) uses this retrieval pathway for returning to the ER. From these results, we conclude that AT(10) can serve as both an ER retention and retrieval signal. By contrast, GFP_ER_- AG(10) did not show apparent Golgi localization even after KDELR2 (Fig. S5), suggesting that AG(10) does not have an ER retrieval function.

### CAT-tails confer differential protein-protein interactions

It is conceivable that different CAT-tails might be recognized by distinct factors either at the ER or in the Golgi, causing different trafficking patterns for TAQC substrates. To test this hypothesis, we transfected HEK293T cells with GFP_ER_-AT(10) or GFP_ER_-AG(10) and affinity-purified these proteins from cell extracts using a GFP antibody. Proteins co-purified were analyzed by SDS-PAGE and silver staining, which suggested that more proteins bound to GFP_ER_-AT(10) than GFP_ER_-AG(10) (Fig. 7A); this was confirmed by mass spectrometry analysis of selected gel sections (Fig. 7B, Supplementary table S2). Analysis of proteins differentially bound to GFP_ER_-AT(10) and GFP_ER_-AG(10) identified two groups of proteins. Group one includes several abundant ER chaperons such as BiP and calnexin (CANX). These proteins showed modest enrichment in GFP_ER_-AT(10) pulldown (Fig. 7C, D). By contrast, proteins in group 2 were enriched in GFP_ER_-AT(10) pulldown by at least 4-fold. This group includes several ERAD factors including Sel1L, a component of the Hrd1 retrotranslocation complex, the Hrd1 cognate conjugating enzyme UBE2J1, and VCP/p97 (Fig. 7C, D). This group also contains a previously uncharacterized Golgi protein GLG1 (Fig. 7D), consistent with the notion that at least a fraction of GFP_ER_-AT(10) has trafficked to the Golgi.

**Figure 7.**
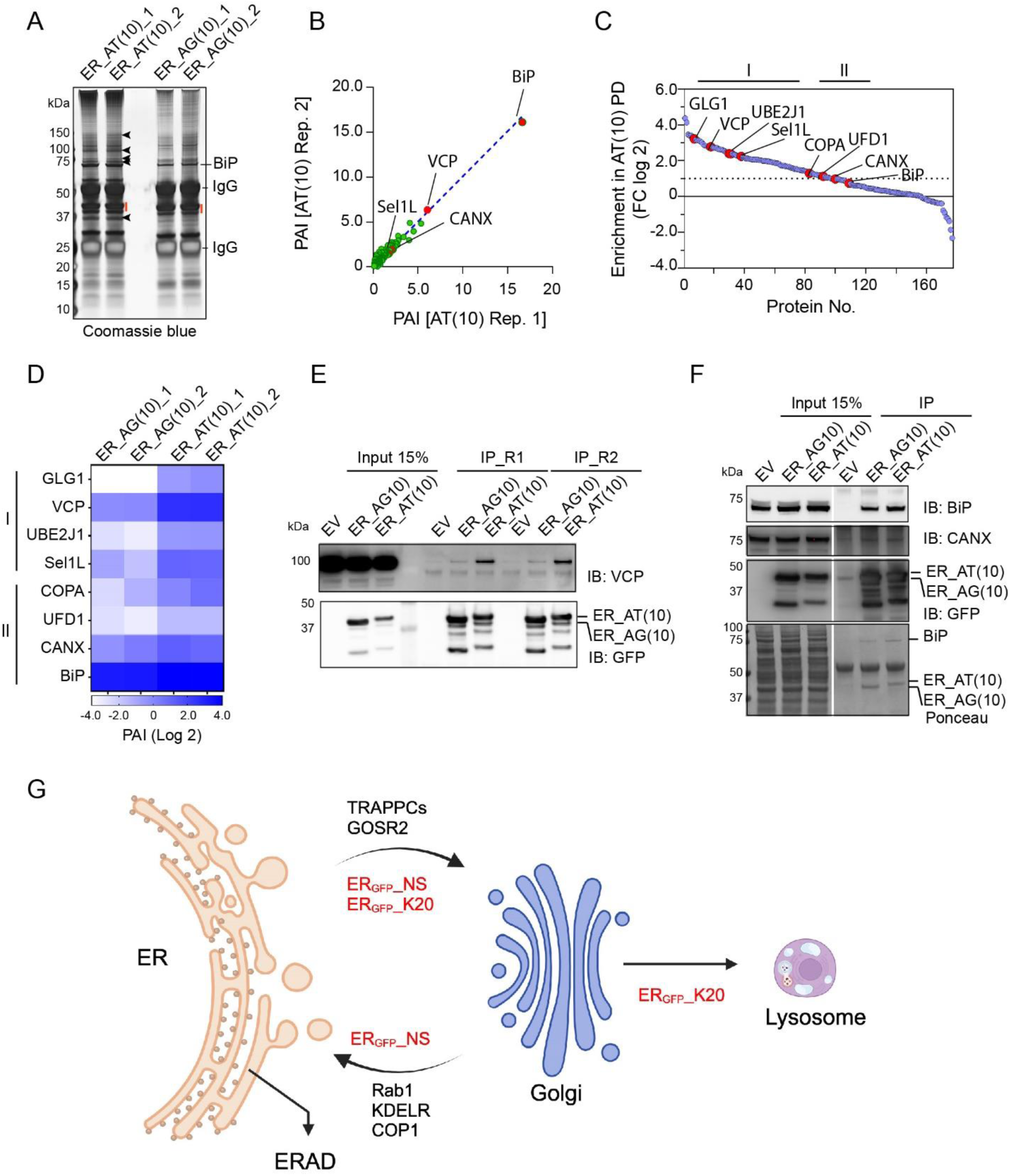
Mass spectrometry analysis of the AT10 and AG10 interactomes. (A) HEK293T cells transfected with the indicated CAT tail mimetics were lysed and subject to affinity purification by GFP antibodies. Purified proteins were analyzed by SDS-PAGE and silver staining. Shown is a result from two biological repeats. (B) A scatter plot shows the relative abundance of proteins co-precipitated with ER_GFP_-AT(10) from two purifications. (C) A scatter plot shows the relative enrichment of the purified proteins in ER_GFP_-AT(10) pulldown (PD). (D) A heat map showing the relative abundance of the indicated proteins in the purification experiments. (E, F) HEK293T cells were transfected with the indicated CAT tail mimetics or an empty vector as a negative control. Cell extracts were subject to immunoprecipitation by FLAG antibodies followed by immunoblotting. R1 and R2 indicated two biological repeats in E. (G) A diagram depicting the distinct branches of TAQC.

We next used immunoblotting to confirm several identified proteins. To exclude non-specific binding, we used cells transfected with an empty vector (EV) as a negative control. Consistent with the mass spectrometry result, BiP and CANX were co-precipitated with GFP_ER_-AT(10) and GFP_ER_-AG(10) with only a slight enrichment in GFP_ER_-AT(10)-precipitated samples. By contrast, p97 showed more pronounced interaction with GFP_ER_-AT(10) than GFP_ER_-AG(10). This observation further validated ERAD as the major degradation pathway for GFP_ER_-AT(10) but not for GFP_ER_-AG(10).

## Discussions

Our study has revealed multiple degradation pathways for defective nascent chains that clog the Sec61 translocon during co-translational translocation, which resolves a major contradiction in the field. We show that “translocon cloggers” can be transported to lysosomes or targeted to the proteasome. In both cases, they use a trafficking mechanism that depends on RPL26 UFMylation, SAYSD1, and TRAPPC proteins to leave the ER. Once arriving at the Golgi, some substrates, as exemplified by ER_GFP__NS, use a KDEL-dependent ER retrieval pathway to return to the ER for elimination by ERAD. Others (e.g. ER_GFP__K20) continue to reach lysosomes (Fig. 7G). Although we cannot exclude that some “cloggers” in an early phase of protein translocation (those with only a small portion inserted into the ER) might “slip” back into the cytoplasm and be degraded by a mechanism akin to cytosolic RQC (Scavone et al., 2023), our results are consistent with the model that defective translocation products are mostly released into the ER lumen (Arakawa et al., 2016), followed by CAT-tail-mediated sorting before degradation.

As a crucial aspect of the cytosolic RQC pathway, CAT-tailing is a unique co-translational modification mediated by Rqc2p in yeast or NEMF in mammals. It prevents the accumulation of toxic, non-functional proteins in the cell. Thus, this process is vital for maintaining cellular homeostasis, particularly under stress conditions that disrupt normal protein synthesis (Howard and Frost, 2021). Despite its importance, the precise function of CAT-tailing in RQC is not fully elucidated, let alone its role in TAQC.

We and others showed that the efficient degradation of model TAQC substrates in mammalian cells requires LTN1, NEMF, and ERAD components such as Hrd1 (Scavone et al., 2023; Wang et al., 2023), consistent with studies reporting the involvement of Ltn1p and Hrd1p in yeast TAQC (Arakawa et al., 2016; Crowder et al., 2015). Mechanistically, our findings suggest that NEMF-mediated CAT-tailing may provide a critical sorting function in TAQC. NEMF-mediated CAT-tailing in mammalian cells can add an array of short peptide tails to stalled nascent chains (Udagawa et al., 2021). Thus, differential tail recognition by ER quality control factors might specify substrate fate in TAQC. In our model, the precise role of LTN1 in TAQC is not obvious since substrates released into the ER lumen cannot be ubiquitinated by LTN1. However, we and others observed a significant down-regulation of NEMF in LTN1 knockdown cells (Lv et al., 2024), suggesting that LTN1’s contribution to TAQC may be limited to NEMF stabilization.

Using a set of “CAT-tail mimetics,” we show that an AT-rich tail can function as both an ER-retention and an ER-retrieval signal. Accordingly, GFP_ER_-AT(10) tagged with an AT-rich tail is degraded predominantly by the proteasome via ERAD. By contrast, an AG-rich tag can direct substrates to lysosomes, albeit not efficiently. Although we do not know the exact CAT-tail sequence on authentic TAQC substrates, the fact that modified ER_GFP__NS and ssGFP-K20 have distinct aggregation propensity mirrored by GFP_ER_ appended with an AT- and AG-rich tag, suggests that ER_GFP__NS is probably modified with an AT-rich tail, while the tail on ssGFP-K20 might have more AG components. Consistent with this view, like GFP_ER_-AT(10), ER_GFP__NS degradation is mediated mainly by ERAD, while ssGFP-K20 behaves more like GFP_ER_-AG(10), as its degradation can be mediated by both lysosomes and the proteasome. Additional methods are needed to determine the exact amino acid sequence appended to TAQC substrates.

CAT tails in yeast are predominantly AT-rich (Kostova et al., 2017), but the sequence of mammalian CAT tails appears more diversified (Udagawa et al., 2021). Interestingly, the UFMylation system and the UFMylation site on RPL26 are both missing in budding yeast. These observations raise the possibility that the UFMylation system might be evolved to cope with the increased complexity of the CAT-tailing system; UFM1 may serve to recruit downstream TAQC components to help sort distinct substrates to different degradation pathways.

Since CAT-tail-mediated peptide synthesis does not requires a mRNA template, it is unclear how a specific sequence is determined in mammalian cells. ER_GFP__K20 and ssGFP-K20 both use a poly(A) sequence to stall ribosomes. Nevertheless, these two proteins have different fates. This implies that the amino acid sequence context in which ribosome stalling occurs might contribute to CAT-tail sequence specification. Moreover, since lysosome- and proteasome-mediated degradation of ssGFP-K20 can occur in distinct cell populations, our model would suggest that CAT-tail sequence specificity may be further influenced at individual cell levels by other unknown factors. Future studies are required to determine the functional scope of CAT-tailing in TAQC and the mechanism by which CAT-tail sequences are specified.

## Materials and Methods

### Chemicals, plasmids, and antibodies

Bafilomycin A1, NMS-873, and MG132 were purchased from LC Laboratories, Xcess Biosciences, and Millipore, respectively. TAK-243 was from Selleck chemicals. Anisomycin was purchase from Sigma. Bortezomib was from Takeda Pharmaceuticals. Cholera toxin subunit B (CF® 594 conjugated) was from Biotium.

The pER_GFP_-K20 and pER_GFP_-NS plasmids were reported previously (Wang et al., 2020). We purchased the plasmid expressing an ER-localized hook protein from Addgene (Ii-Str_Neomycin; #65308). To generate various model substates with artificial CAT-tails, we used restriction enzymes BamHI and XbaI to cut pER_GFP_-K20 (Addgene plasmid #133861), removing most sequence downstream GFP-3xFLAG. Various CAT-tail encoding sequences (listed below) were synthesized by IDT and PCR-amplified using the corresponding primer pairs. PCR products were digested with BamH1 and XbaI. The digested products were ligated into digested pER_GFP_-K20. To make pGFP_ER_-ext, a stop codon was introduced at the site to include a short nucleotide (encoding NH2-AGSPGPTPSGTNVGS-COOH) in the ORF downstream of 3xFLAG by site-directed mutagenesis. To make pGFP_ER,_ a stop codon was introduced right after 3xFLAG-conding sequence.

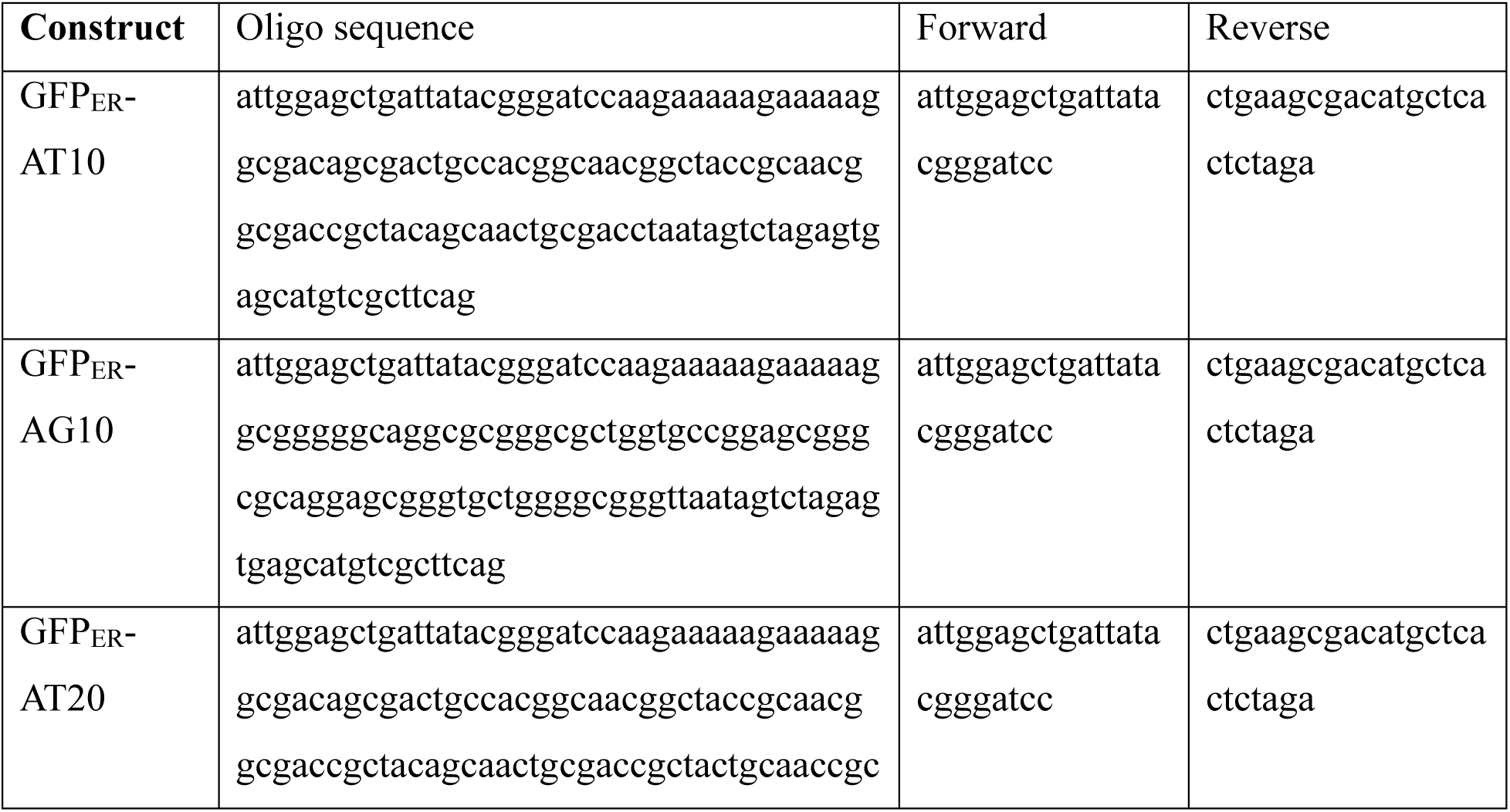
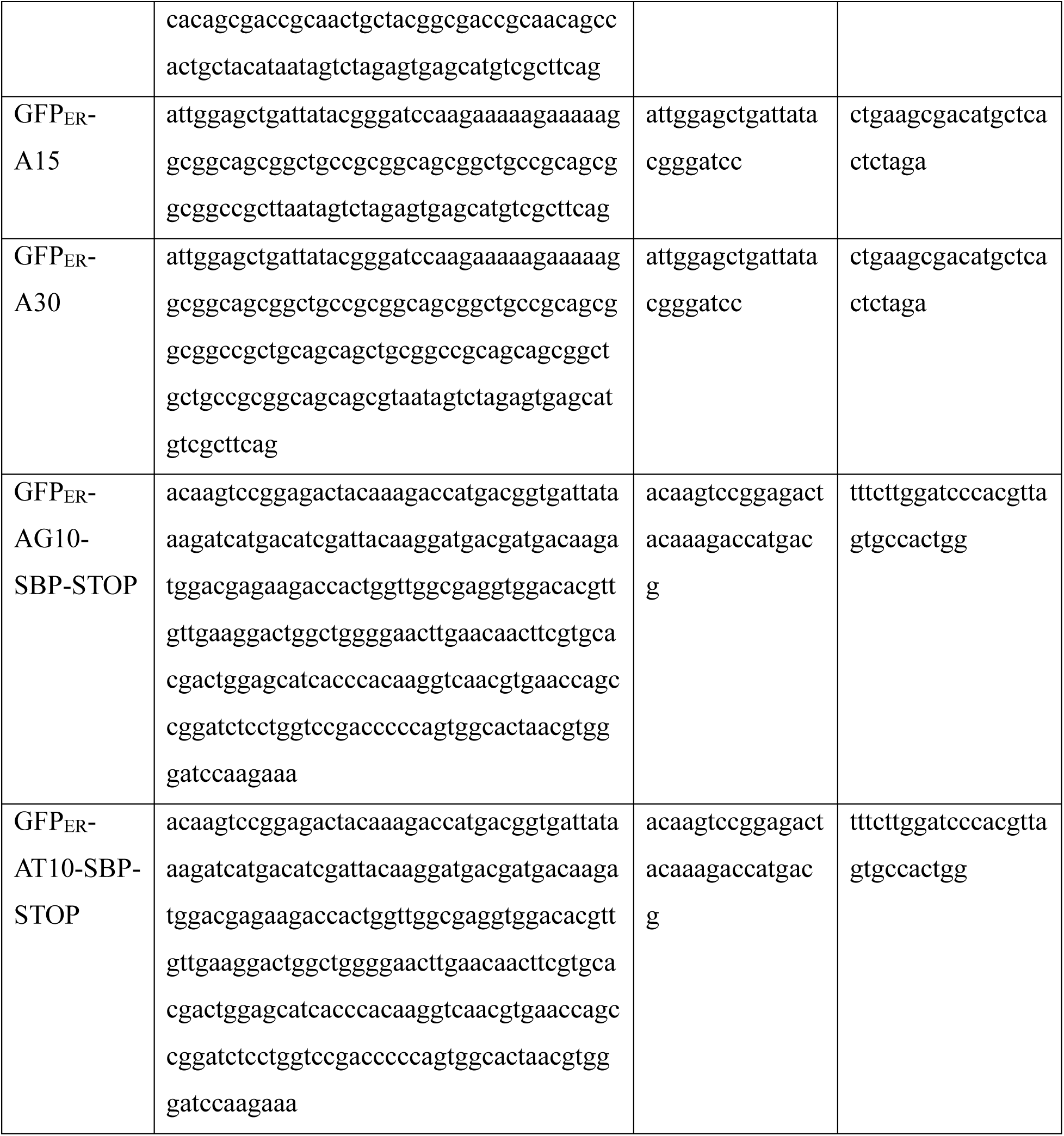

### Inhibitor treatments

To test the effect of several inhibitors on the expression and trafficking of _SS_GFP-K20 and model substrates with artificial CAT-tails, U2OS were seeded at 25,000 cells/well on an 8-well chamber. After 24 h, cells were co-transfected with 250 ng _SS_GFP-K20 plus 50 ng mCh-Lamp1 (to identify localization of _SS_GFP_K20 expression) or with 200 ng CAT-tail-containing model substrates using Invitrogen’s Lipofectamine 2000 transfection reagent. Twenty-four hours later, cells were treated for ∼6 h with either 200 nM Baf A1, 20 µM MG132, or 1 µM Bortezomib, and imaged by live cell imaging. To cause partial translation arrest, cells were treated with 200 nM anisomycin for 1 h before analysis. To test the degradation pathway for _SS_GFP_K20, cells transfected with _SS_GFP_K20 for 24 h were treated with 200 nM Baf A1, 20 µM MG132, 1 µM Bortezomib, 5 µM NMS-873, or 10 µM TAK243 before imaging.

### Genome-wide CRISPR/Cas9 knockout screen

Genome-wide CRISPR/Cas9 knockout screen was performed as described previously (Wang et al., 2023). Briefly, the GeckoV2 library (Addgene #1000000048) was amplified according to the online protocol (Sanjana et al., 2014) and the complexity of the library determined by high-throughput sequencing. Lentiviruses expressing the sgRNA library and Cas9 were used for spin-infection as follows: 60 million ER_GFP__NS-expressing stable HEK293T cells supplemented with 8 μg/mL polybrene (Sigma) were seeded into two 12-well plates at a density of 3 million cells/well. Concentrated GeCKO v2 lentivirus was added to each well at a multiplicity of infection (MOI) of 0.3. Cells were spun at 1,000 g at room temperature for 2 h followed by incubation at 37 °C in a humidified incubator for 1 h. After medium removal, fresh medium was added and cells were incubated for 48 h before puromycin selection (0.3 μg/mL). The transduced cells were subcultured in medium supplemented with puromycin every 2 days for a total of 8 days and 80 million cells were maintained for each passage. After puromycin treatment, cells were recovered in a medium lacking puromycin for 24h before sorting. In total, 60 million cells were sorted into GFP high (1% of total cells) or GFP low (80% of total cells) cell populations by a FACS AriaII cell sorter. Genomic DNA was extracted from each cell population using a QIAGEN Blood Maxi kit (for GFP low cells) or QIAGEN Blood Midi kit (for GFP high cells) according to the manufacturer’s instructions. sgRNAs were amplified from genomic DNA samples using the Herculase II Fusion DNA polymerase (Agilent Technologies) in two PCR steps as follows. In the first PCR, the genomic region containing sgRNAs were amplified from 130 μg total DNA using the following primers: Forward: 5′-AATGGACTATCATATGCTTACCGTAACTTGAAAGTATTTCG-3′; Reverse: 5′-CTTTAGTTTGTATGTCTGTTGCTATTATGTCTACTATTCTTTCC-3′. In total, 13 PCR reactions were performed in parallel, with 10 μg genomic DNA in each reaction using Herculase II Fusion DNA Polymerase (Agilent) for 18 cycles and the resulting PCR products were combined. In the second step PCR, 5 μL of the first PCR product was used in a 100 μL reaction volume and 24 PCR cycles was used. The primers used for the second PCR include stagger sequences of variable lengths and a 6-bp barcode for multiplexing of different biological samples. The second step PCR primers used are: Forward: 5′-AATGATACGGCGACCACCGAGATCTACACTCTTTCCCTACACGACGCTCTTCCGATCT (1-9bp variable length sequence)-(6bp barcode)tcttgtggaaaggacgaaacaccg-3’; Reverse: 5′-CAAGCAGAAGACGGCATACGAGATGTGACTGGAGTTCAGACGTGTGCTCTTCCGATCTtctactattctttcccctgcactgt-3′. The PCR products were gel extracted, quantified, and sequenced using a NovaSeq sequencer (Illumina) by the NHLBI DNA Sequencing and Genomics Core. sgRNA sequences were obtained per sample by extracting 20 bps followed by the index sequence of “TTGTGGAAAGGACGAAACACCG” on the de-multiplexed FASTQ files from Illumina’s NGS sequencer using the Cutadapt software, version 2.8 (https://doi.org/10.14806/ej.17.1.200). FASTQC, version 0.11.9 was used to assess the sequencing quality (http://www.bioinformatics.babraham.ac.uk/projects/fastqc/). MAGeCK, version 0.5.9, was used to quantify and to identify differentially expressed sgRNAs. MAGeCK count command was run on the merged two-half libraries of A and B to quantify. Differentially expressed sgRNAs with statistical significance were determined by running the MAGeCK test command in paired mode. Genes were ranked based on the number of unique sgRNA enriched in the GFP high population versus the GFP low population. Data was derived from two biological repeats.

### Cells and siRNA transfections

For all experiments, HEK293T and U2OS cells (ATCC) were cultured in Dulbecco’s Modified Eagle Medium (DMEM, Corning) containing 10 % fetal bovine serum (FBS) and antibiotics (Penicillin/streptomycin, 10 U/mL) at 37°C in a 5 % CO2 humidified incubator. To test the effect of NEMF and UFM1 knockdowns on ER_GFP__NS expression by microscopy, U2OS cells were seeded at 25,000 cells/well on an 8-well chamber (Ibidi). Twenty-four h after seeding, cells were transfected with 4 pmol of siRNAs targeting genes specified in the figure legends using Invitrogen RNAiMax transfection reagent. 24 h later, cells were transfected with 300 ng ER_GFP__NS with Invitrogen Lipofectamine 2000 transfection reagent. Cells were incubated with fresh media for 24 h before fixation and imaging. A similar protocol was used to knock down COPB1, KDELR2, RAB1B and TMED3 in HEK293T cells to check the role of these genes in ER_GFP__NS trafficking by fluorescence microscopy. In some experiments, a Cy3 labeled negative control siRNA was included to verify transfection efficiency.

To test the effect of NEMF and UFM1 knockdowns on ER_GFP__NS protein level by immunoblotting, HEK293T cells were seeded on a 6-well plate at 500,000 cells/well. Plates were pre-coated with Poly-D-lysine to assist with cell adherence to the well. Twenty-four hours after seeding, cells were transfected with 120 pmol of siRNAs (a 1:1:1 mixture of 3 siRNAs targeting the same gene) using Invitrogen RNAiMax transfection reagent. After an additional 24 h, cells were split 1:2 onto pre-coated 12-well plates and 4 h later were transfected with 500 ng ER_GFP__NS using *Trans*IT-293 Transfection Reagent (Mirus). Media was changed after 24 h, then cells were incubated for an additional ∼20 h before cell extracts were made with NP40 lysis buffer (0.5 % NP40, 50 mM Tris-HCl pH 7.4, 2 mM MgCl_2_, 150 mM NaCl, 1 mM EDTA) plus 1000x protease inhibitor and 0.5mM TCEP. Cells were washed with cold 1x PBS, then 150µL NP40 lysis buffer was added directly to wells and rocked for 30 min at 4 °C. After lysis, lysate was transferred to a microcentrifuge tube and centrifuged at top speed for 5 min. The supernatant (120 µL) was collected in a clean microcentrifuge tube and 40 µL of 4x Laemmli sample buffer (plus β-mercaptoethanol) was added to each tube. Samples were heated at 90°C for 15min and loaded on an SDS-PAGE gel for immunoblotting analysis. siRNAs targeting UFM1, SAYSD1, and various TRAPPC genes were reported previously(Wang et al., 2023). Other siRNAs are listed below.

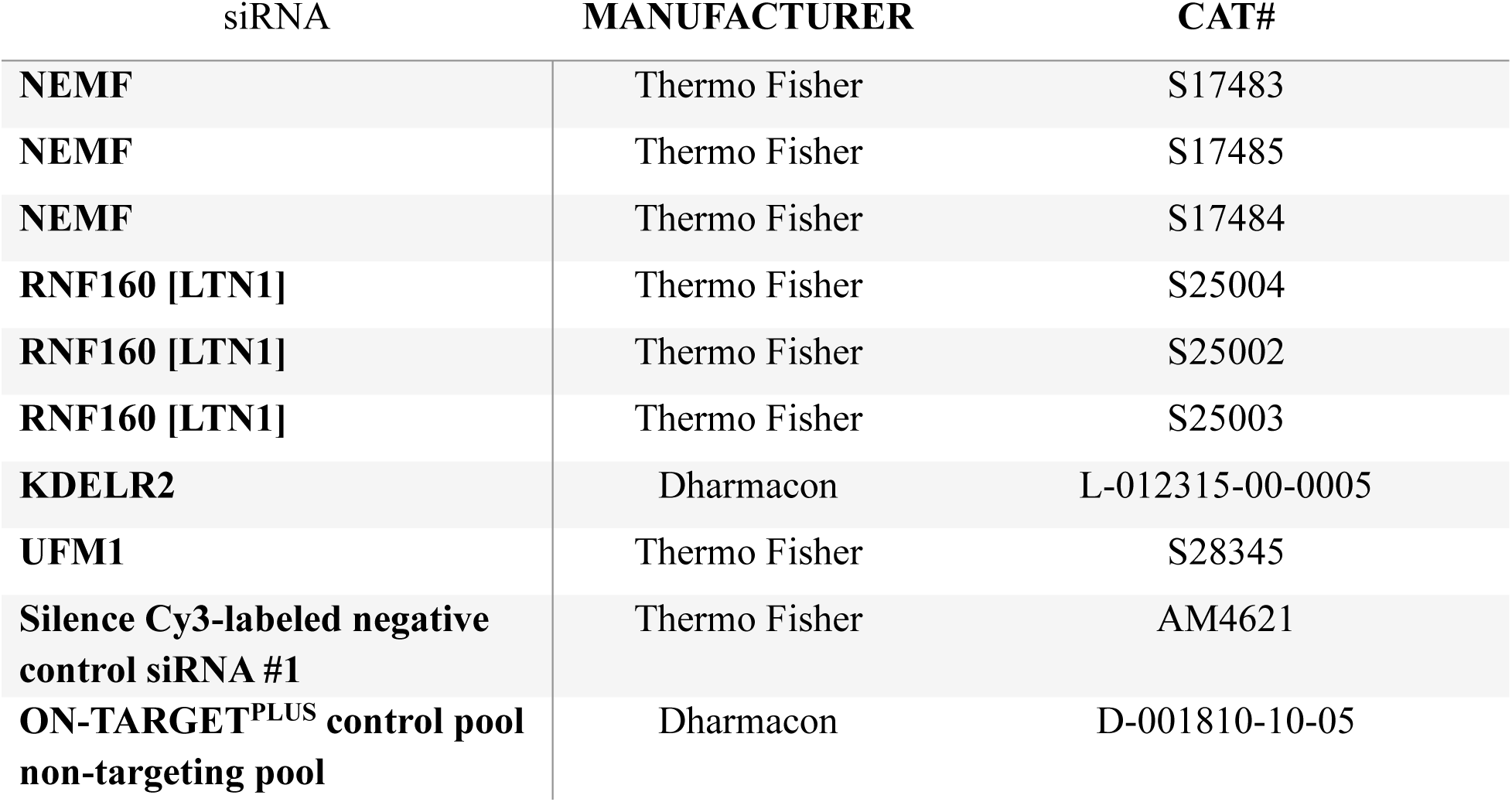
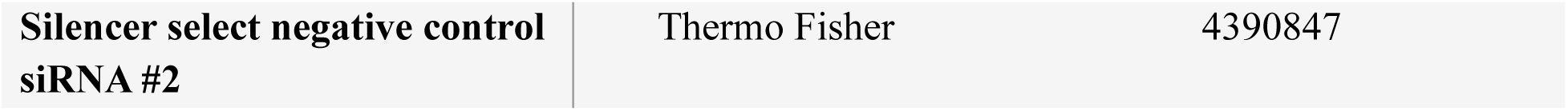

### RUSH Assay, live cell imaging, and image analyses

To explore trafficking of GFP_ER_-SBP-AG10 (the bait), U2OS cells were seeded at 25,000 cells/well on an 8-well chamber. Twenty-four h after seeding, cells were co-transfected with 75 ng ER_GFP-SBP-AG10 and either 225 ng Ii-Str_Neomycin (A gift from F. Perez, Addgene plasmid #65308; http://n2t.net/addgene:65308; RRID:Addgene_65308) or EV using Invitrogen Lipofectamine 2000 transfection reagent. After an additional 24 h, media was replaced with fresh media. Forty-eight h post-transfection, cells were treated for ∼4-5 h with DMSO and 80 µM D-biotin as indicated and imaged with live cell imaging. Cells were imaged using a Nikon C1 SORA spinning disk confocal microscopy. Images were processed by Image J or Imaris for 3D visualization. Fluorescence intensity was measured by Image J.

### Co-immunoprecipitation, affinity purification and mass spectrometry

To test the interaction of TAQC substrates with Sec61β, ribosome, and SAYSD1, HEK293T cells were seeded in a 6-well plate at 0.5 million/well and grown for 24 h followed by transfection of 1.5 μg ER_GFP__NS-expressing plasmid. For control experiments, cells were transfected with a pcDNA3 empty vector. Forty-eight h post-transfection, cells were washed with ice-cold PBS and lysed in 0.5 mL CHAPS lysis buffer (1% CHAPS, 50 mM HEPES pH 7.3, 100 mM NaCl, 1 mM DTT, and complete protease inhibitors) for 15 min at 4 °C. The cell extracts were then centrifuged at 15.6 kg for 10 min at 4 °C. To immunoprecipitate ER_GFP__NS, cleared supernatants were incubated with 30 μL protein A beads coated with GFP antibodies at 4°C for 1 h using a head-over-tail rotator. Precipitated samples were washed with CHAPS lysis buffer two times before SDS-PAGE analyses.

For large scale affinity purification of GFP_ER_-AT(10) and GFP_ER_-AG(10), HEK293T cells grown in four 10cm dishes were transfected with 6 µg CAT-tail mimetic DNA using TransIT293 (Mirus Bio). Cells were expanded for 72 h and then harvested in ice cold PBS. After centrifugation, cell pellets were lysed in a buffer (4.5 mL/plate) containing 1% CHAPS, 50 mM HEPES pH 7.3, 100 mM NaCl, 2 mM MgCl_2_, 2 mM CaCl_2_, 1 mM DTT and complete protease inhibitors. Cleared cell extracts were incubated with 200 µL Protein A beads pre-incubated with 10 µg affinity purified GFP antibodies at 4 °C for 60 min. The beads were thoroughly washed with a CHAPS wash buffer containing 0.2 % CHAPS, 50 mM HEPES pH 7.3, 100 mM NaCl, 2 mM MgCl_2_, 2 mM CaCl_2_, 1 mM DTT and complete protease inhibitors. Bound proteins were eluted by Laemmli sample buffer and analyzed by SDS-PAGE gel, followed by silver staining or Coomassie blue staining.

To identify proteins in pulldown experiments, gel sections of interest were first cut to small pieces and washed three times with 25 mM ammonium biocarbonate (NH_4_HCO_3_) in 50% acetonitrile (ACN), and then dried by Speedvac and rehydrated in a 25 mM NH_4_HCO_3_ solution containing trypsin. The amount of trypsin used depended on the amount of protein loaded on the gel. The protein gel pieces were digested overnight at 37 °C. Peptides were extracted by high performance liquid chromatography (HPLC) grade water followed by three washes with 50% ACN/5% formic acid (FA) at room temperature. The combined supernatants were dried down by Speedvac and redissolved in 0.1% FA for further cleaning by C18 tips (Agilent) and dried. The cleaned tryptic digest peptides were dissolved in a buffer with 2% FA and 3% ACN for mass spectrometry analysis.

The digested peptide were subjected to LC MS/MS analysis using an UltiMate 3000 RSLC system (Thermo Fisher Scientific) coupled in-line to an Orbitrap Fusion Lumos mass spectrometer (Thermo Fisher Scientific). Reverse-phase separation was performed on a 50 cm × 75 μm I.D. Acclaim® PepMap RSLC column. Peptides were eluted using a gradient of 4% to 22% B over 87 minutes at a flow rate of 300 nL min−1 (solvent A: 100% H_2_O, 0.1% formic acid; solvent B: 100% acetonitrile, 0.1% formic acid). Each cycle consisted of one full Fourier transform scan mass spectrum (375–1500 m/z, resolution of 120 000 at m/z 400) followed by data-dependent MS/MS scans acquired in the Orbitrap with HCD NCE 30% at top speed for 3 seconds. Target ions already selected for MS/MS were dynamically excluded for 30 s. LC MS/MS data were extracted using MSConvert (ProteoWizard 3.0.10738) and subjected to protein prospector (UCSF) for database searching against a concatenated database consisting of the normal and random form of the human protein database. Trypsin was set as the enzyme with a maximum of two missed cleavage sites. The mass tolerance for parent ion was set as ±20 parts per million, whereas ±0.6-Da tolerance was chosen for the fragment ions. Chemical modifications such as protein N-terminal acetylation, methionine oxidation, N-terminal pyroglutamine, and deamidation of asparagine were selected as variable modifications during database search. The search compare program in protein prospector was used for summarization, validation, and comparison of results.

The relative abundance of the identified proteins in mass spectrometry experiment was calculated using the following function: 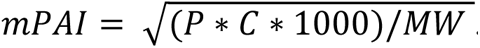 . mPAI (modified Protein Abundance Index), P = total peptide count, C = Sequence coverage, MW = Molecular Weight (kDa). This function is based on previously defined PAI (Ishihama et al., 2005) by incorporating protein sequence coverage as another indicator of abundance, assuming a linear correlation between protein abundance and sequence coverage. To compare protein abundance between AG(10) and AT(10) pulldowns, we assumed a similar peptide count and coverage as the lowest detected protein in the same sample for undetected proteins.

### Immunostaining and immunoblotting

To prepare cells for imaging, cells grown and transfected in a Ibidi imaging chamber were washed once with ice cold phosphate buffer saline (PBS), fixed with 4% paraformaldehyde in PBS for 15-20 min at room temperature, then washed 3 times with PBS. Cells were treated with PBS containing 0.1% NP40 and 5% fetal bovine serum for 15 min at room temperature and then stained for 1 h at room temperature with the primary antibodies listed below. Stained cells were washed 3 times with PBS and then stained with the corresponding secondary antibody conjugated a fluorescence dye (Alexa^488^, Alexa^566^, or Alexa^640^). In some experiments, cells were also stained with Hoechst 33342 to reveal the nuclei. Stained cells were washed 3 times with PBS. Images were acquired by a Nikon CSUW1 -SoRa spinning disc confocal equipped with a 60 x TIRF lens. For immunoblotting analyses, cell lysates or immunoprecipitated materials were fractionated on 4-12% NuPAGE gels following manufacture’s instruction (ThermoFisher). Proteins were transferred to nitrocellular membranes (BioRad) using a semi-dry protein transfer apparatus (Trans-Blot Turbo, BioRad). Membranes were treated with PBS containing 5% non-fat milk to block non-specific binding and then incubated with primary antibodies in PBS containing 5% BSA as indicated below. After washing with PBS three times, we incubated the membranes for 1 h with HRP-conjugated or fluorescent secondary antibodies (Invitrogen). Chemiluminescent or fluorescent signals were detected by the BioRad ChemDoc scanning system.

| **Associated figures** | **Assay type** | **Protein name** | **Manufacturer** | **Catalogue No.** | **Dilution** |
| --- | --- | --- | --- | --- | --- |
| 1D | IF | Nucleolin | Bethyl | A300-711A | 1:1000 |
| 1J | IB | UBE1 | Abcam | 10R-P104A | 1:1000 |
| 1J | IB | Sec61β | A gift from Tom Rapoport lab | | 1:5000 |
| 1J, I, 2C,<br>D, 4D, 5B,<br>7E, F | IB | GFP | Santa Cruz | SC-9996 | 1:500 |
| 1I, 2C, 4D | IB | UFM1 | Abcam | Ab109305 | 1:1000 |
| 1I, 3D, 5B,<br>7E | IB | VCP/p97 | Fitzgerald | 10R-P104A | 1:2000 |
| 2C, D | IB | SAYSD1 | Abcam | Ab189918 | 1:1000 |
| 2D | IB | RPL26 | Bethyl | A300-686A | 1:1000 |
| 3D | IB | Hrd1 | Generated in house<br>(Zhang et al.,<br>2015) |  | 1:1000 |
| 4A | IF | GFP | Thermo Fisher | A11122 | 1:200 |
| 4C | IB | NEMF/SDCCAG1 | Proteintech | 11840-1-AP | 1:1000 |
| 4C | IB | LTN1/RNF160 | Proteintech | 28452-1-AP | 1:2000 |
| 4C | IB | HSP90 | Santa Cruz | SC-13119 | 1:1000 |
| 6C | IF | LAMP1 | Santa Cruz | SC-18821 | 1:500 |
| 6E | IF | GM130 | BD Biosciences | 610822 | 1:500 |
| 7F | IB | BiP | R&D Systems | MAB4846 | 1:1000 |
| 7F | IB | CANX | Enzo | SPA-865 | 1:1000 |

### Statistical analysis

Immunoblot data was analyzed with BioRad Image Lab. Imaging data was analyzed with ImageJ, Imaris, or Nikon Elements Analysis (live cell only). All statistical analyses were completed with GraphPad Prism 10 and are displayed as mean ± s.e.m.

## Acknowledgements

We thank the Advanced Light Microscope Core at NIDDK for assistance with imaging, NHLBI flow cytometry core for cell sorting, and NHLBI Genomic Core for high-throughput sequence. The research is supported by an intramural research program of NIDDK and by NIH grant R35GM145249 (L. Huang).

## Author contributions

A. Ennis, L. Wang, Y. Xu, L. Saidi, and Y. Ye performed the experiments and analyzed the data. X. Wang, C. Yu, and L. Huang designed and performed the mass spectrometry study, S. Yun analyzed the CRISPR screen data. A. Ennis and Y. Ye wrote the paper. All authors participated in editing the manuscript.

