## Supplemental figures and legends for "NEMF-mediated CAT-tailing defines distinct branches of translocation-associated quality control"

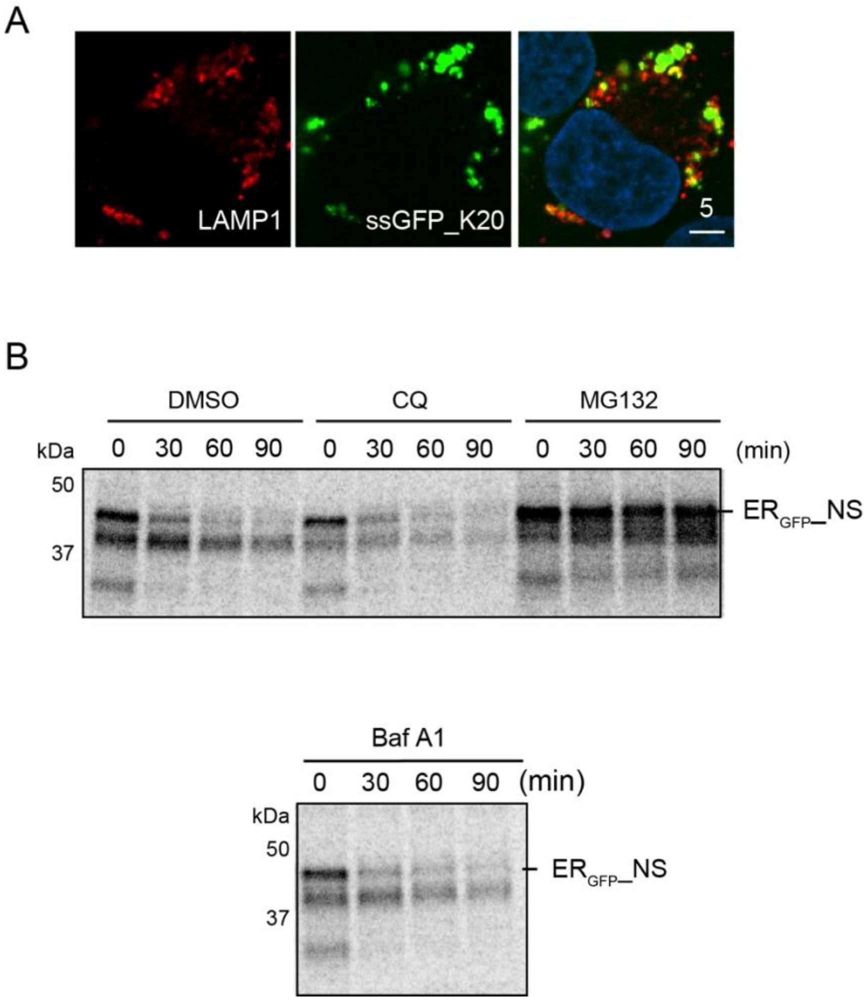


**Figure S1 TAQC substrates can be degraded by lysosomes and the proteasome.**

(A) $_{ss}$GFP_K20-transfected cells were treated with Baf A1 (200 nM 4 h) and stained with LAMP1 antibodies (red). Note the partial localization of $_{ss}$GFP_K20 with LAMP1. Scale bar, 5 μm. (B) ER$_{GFP}$_NS-expressing HEK293T cells treated with the indicated compounds were pulse-labeled with S$^{35}$-Met and chase in a medium containing unlabeled S$^{35}$-Met. Samples taken at the indicated time points were subject to immunoprecipitation and autoradiography. CQ, Chloroquine.

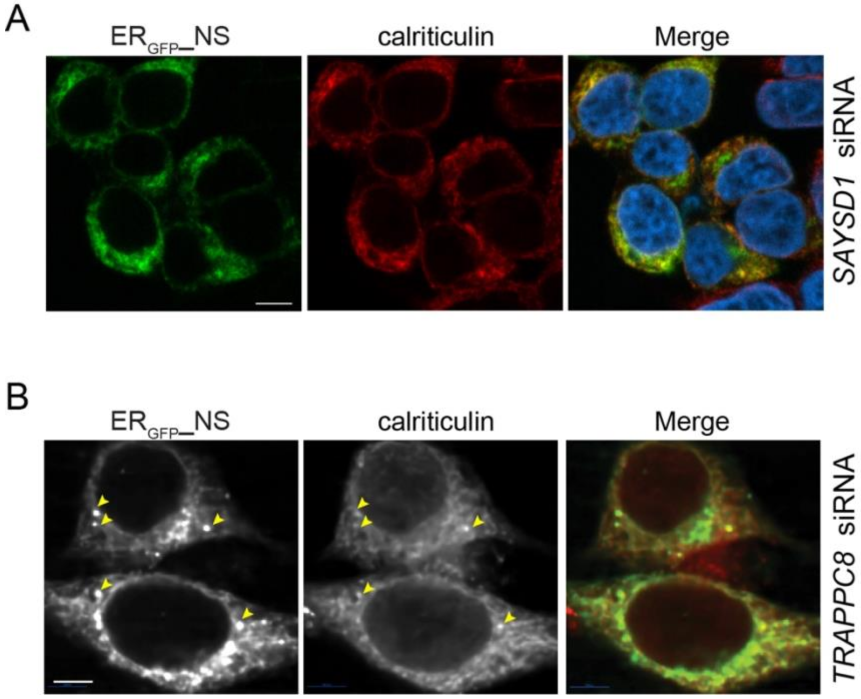


**Figure S2 ER$_{GFP}$_NS accumulates in the ER in SAYSD1 and TRAPPC8 knockdown cells.** (A) HEK293T cells transfected with ER$_{GFP}$_NS and *SAYSD1* siRNA were stained with calreticulin antibodies (red) and Hoechst (blue). Scale bar, 5 μm. (B) HEK293T cells transfected with ER$_{GFP}$_NS and *SAYSD1* siRNA were stained with calreticulin antibodies. Arrows indicate a few example of ER$_{GFP}$_NS and calreticulin colocalization. Scale bar, 5 μm.

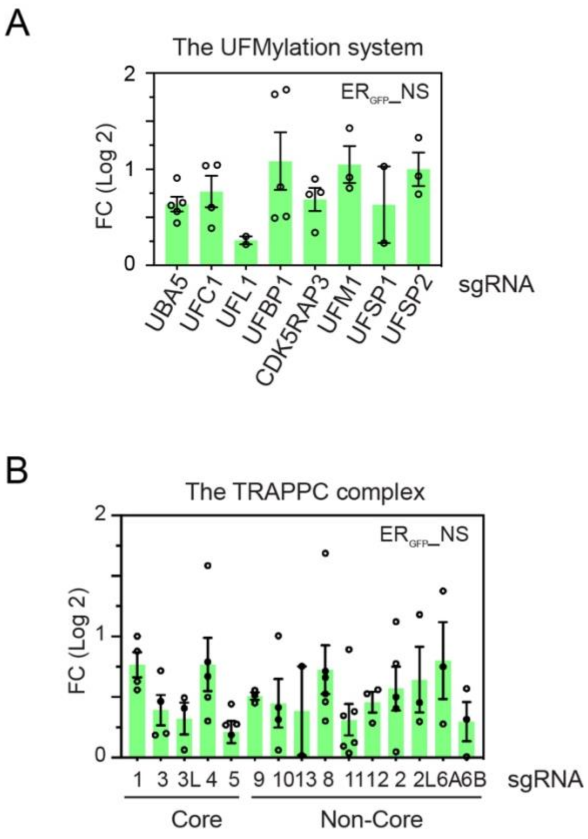


**Figure S3 Genetic screen reveals the UFMylation system and the TRAPPC complex as regulators of ER$_{GFP}$_NS degradation.**

sgRNAs targeting genes of the UFMylation system (A) or the TRAPPC complex (B) are positively enriched in GFP high population, suggesting they are positive regulators for ER$_{GFP}$_NS degradation.

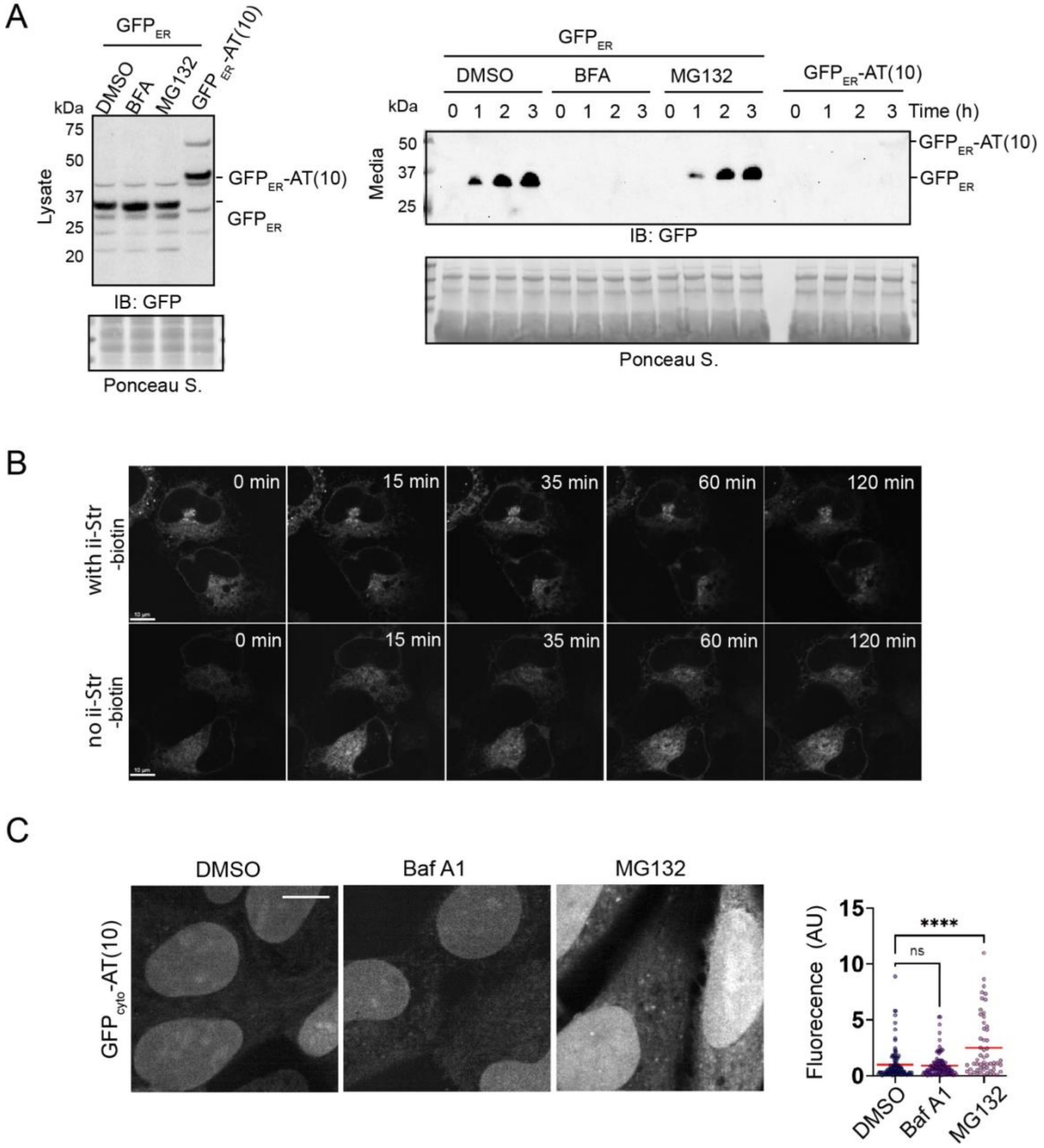


Figure S4 The AT(10) tail inhibits GFP$_{ER}$ secretion but can still supports the transport of GFP$_{ER}$ to the Golgi.

(A) Conditioned media and lysates from HEK293T cells transfected and treated as indicated were analyzed by immunoblotting. (B) U2OS cells were transfected with GFP$_{ER}$-SBP-AG(10) either with or without the ER trap ii-Str. Cells were treated with biotin (top panels) or untreated (bottom panels) for the indicated time points. (C) Cells transfected with GFP$_{cyto}$-AT(10) as a cytosolic CAT-tailing mimetics were treated with the indicated compounds for 6 h and then imaged and quantified. AU, arbitrary unit. By One-way ANOVA's multiple comparison. ns, Non-significant. ****, p<0.0001. Scale bars, 10

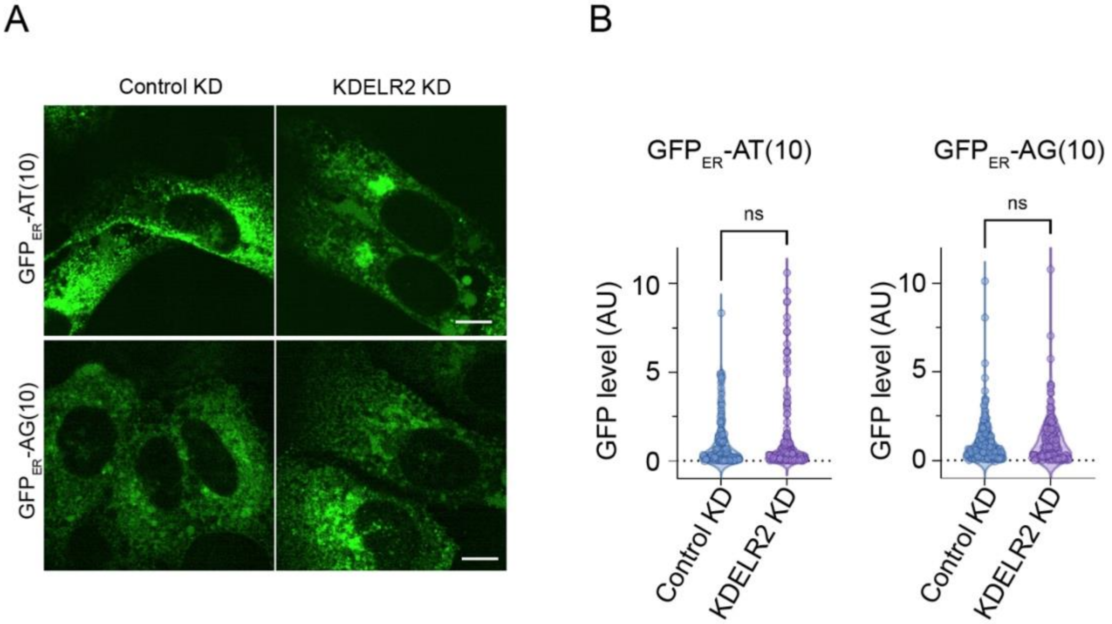


Figure S5 KDELR2 knockdown increases GFPER-AT(10) in the Golgi.

(A) U2OS cells transfect with a control or KDELR2 siRNA and the indicated ER CAT-tailing mimetics were imaged. Scale bars, 10 μm. AU, arbitrary unit.

**Movie 1 A RUSH assay reveals the ER-to-Golgi trafficking of $GFP_{ER}$-SBP-AG(10).**
U2OS cells transfected with $GFP_{ER}$-SBP-AG(10) and an ER localized hook ii-str were treated with biotin (80 μM) and then imaged by live cell confocal microscopy.

**Movie 2 $GFP_{ER}$-AT(10) is co-localized with GM130 in U2OS cells.**
U2OS cells transfected with $GFP_{ER}$-AT(10) were fixed and stained with GM130 antibody and then imaged in 3D by confocal microscopy. Shown is a reconstructed rotational 3D views by Imaris.

**Table S1 A list of sgRNAs enriched in GFP high cells in $ER_{GFP}$_NS screen.**

**Table S2 A list of proteins identified as potential interactors of $GFP_{ER}$-AT(10).**
